# Variation and inheritance of small RNAs in maize inbreds and F_1_ hybrids

**DOI:** 10.1101/692400

**Authors:** Peter A. Crisp, Reza Hammond, Peng Zhou, Brieanne Vaillancourt, Anna Lipzen, Chris Daum, Kerrie Barry, Natalia de Leon, C. Robin Buell, Shawn M. Kaeppler, Blake C. Meyers, Candice N. Hirsch, Nathan M. Springer

**Author notes:** **Author for contact details:** Nathan Springer. **Author Contributions:** N.S., C.H., B.M., S.K., C.R.B., N.dL. conceived the experiment. B.V., A.L., C.D., K.B. performed the experiments. P.C., R.H., P.Z. analyzed the data. P.C. and N.S. wrote the manuscript with contributions from all authors. All authors have read and approved the manuscript. N.S. agrees to serve as the author responsible for contact and ensures communication.

## Abstract

Small RNAs (sRNAs) regulate gene expression, play important roles in epigenetic pathways, and have been hypothesised to contribute to hybrid vigor in plants. Prior investigations have provided valuable insights into associations between sRNAs and heterosis, often using a single hybrid genotype or tissue. However, our understanding of the role of sRNAs and their potential value to plant breeding are limited by an incomplete picture of sRNA variation between diverse genotypes and development stages. Here, we provide a deep exploration of sRNA variation and inheritance among a panel of 108 maize samples spanning five tissues from eight inbred parents and 12 hybrid genotypes, covering a spectrum of heterotic groups, genetic variation, and levels of heterosis for various traits. We document substantial developmental and genotypic influences on sRNA expression, with varying patterns for 21-nt, 22-nt and 24-nt sRNAs. We provide a detailed view of the distribution of sRNAs in the maize genome, revealing a complex make-up that also shows developmental plasticity, particularly for 22-nt sRNAs. sRNAs exhibited substantially more variation between inbreds as compared to observed variation for gene expression. In hybrids, we identify locus-specific examples of non-additive inheritance, mostly characterised as partial or complete dominance, but rarely outside the parental range. However, the global abundance of 21-nt, 22-nt and 24-nt sRNAs varies very little between inbreds and hybrids, suggesting that hybridization affects sRNA expression principally at specific loci rather than on a global scale. This study provides a valuable resource for understanding the potential role of sRNAs in hybrid vigor.

**One-sentence summary:** Characterizing the roles of development and genotype in driving expression variation of different small RNA populations in maize inbreds and their F_1_ hybrids.

## Introduction

Molecular variation is widely studied to understand the basis of plant traits. This can include variation in DNA sequence, DNA methylation or chromatin modifications, changes in abundance of messenger RNA (mRNA) or small RNAs (sRNAs), as well as changes in the proteome or metabolome. These types of variation differ substantially in their heritability across generations and stability in different cells or tissues of the same organism. Changes in DNA sequence are highly stable among the cells of an organism while the abundance of mRNAs or metabolites can vary during development or in response to environmental conditions. This stability is a key factor when considering experimental designs for detecting molecular variation and for attempting to link molecular variation to plant traits. In this study, we monitored variation in sRNA abundance in different tissues and genotypes of maize to understand the patterns of sRNA variation and inheritance.

Heterosis, or hybrid vigor, occurs when an F_1_ hybrid outperforms either parent. The contribution of various potential molecular mechanisms to heterosis remains one of the most intriguing and powerful enigmas in plant biology (Hochholdinger and Baldauf, 2018; Birchler et al., 2003). While the molecular basis of heterosis remains uncertain, it is clear that variation between members of the same species is a requirement (Schnable and Springer, 2013). This variation can occur at the genomic, transcriptomic and epigenetic level (Chen, 2013). Unfortunately, we still lack a coherent molecular mechanism to explain the sources of non-additive inheritance patterns of gene expression in hybrids (Birchler et al., 2010). However, a variety of studies have found evidence that epigenetic factors including sRNAs and DNA methylation play a role in heterosis [reviewed in (Greaves et al., 2015; Ryder et al., 2014; Groszmann et al., 2013)]. Small RNAs are promising candidates for modulating the epigenome and gene expression in hybrids. sRNAs can regulate gene expression via post-transcriptional gene silencing and through transcriptional silencing by directing changes in DNA methylation (Borges and Martienssen, 2015). The canonical functional size classes of endogenous sRNAs in plants include 21-nucleotide (nt), 22-nt and 24-nt sRNAs (Axtell, 2013). 21-nt sRNAs are mainly composed of microRNAs (miRNAs) that regulate the expression of mRNAs, while the 24-nt size class is predominantly produced by the RNA-directed DNA Methylation (RdDM) pathway to control transposable elements (TEs) (Borges and Martienssen, 2015). In maize, RdDM activity is localized to gene flanks (Gent et al., 2013; Li et al., 2015a), where 24-nt sRNAs are abundant (Xin et al., 2014; Wang et al., 2009) and positively correlate with expression levels of the flanking gene (Lunardon et al., 2016; Gent et al., 2013). In fact, the bulk of the heterochromatic maize genome may be incompatible with small RNA production (Gent et al., 2014). Among the species that have been profiled, the 22-nt size class stands out in maize because of its abundance relative to the other classes of sRNAs. A detailed analysis of a 22 Mb contiguous region of the maize genome found that 22-nt sRNAs are enriched in the highly repetitive regions (Wei et al., 2009; Nobuta et al., 2008; Regulski et al., 2013). Despite our knowledge of sRNA in many species, including maize, there are open questions about the sources of sRNA variation among maize lines, the inheritance pattern of this variation, and how it might contribute to heterosis.

Prior studies have compared sRNAs profiles between hybrids and their parents across various species, including in Arabidopsis (Groszmann et al., 2011; Shen et al., 2012; Li et al., 2012, 2015b), rice (Chen et al., 2010; Chodavarapu et al., 2012; He et al., 2010), tomato (Shivaprasad et al., 2012), wheat (Kenan-Eichler et al., 2011) and maize (Barber et al., 2012; He et al., 2013; Regulski et al., 2013; Seifert et al., 2018a, 2018b; Xin et al., 2014). Collectively, these investigations indicate that like mRNA expression, sRNA expression can be inherited non-additively. sRNAs tend to have reduced expression in hybrids relative to parents, particularly 24-nt sRNAs (Groszmann et al., 2011), and this trend is most evident where sRNAs are differentially expressed between parents (Groszmann et al., 2011; Shen et al., 2012; Barber et al., 2012). sRNAs are also associated with changes in DNA methylation and changes in gene expression in hybrids (Chodavarapu et al., 2012; Greaves et al., 2014; Shen et al., 2012; Li et al., 2015b). In maize, associations between sRNA expression and heterotic traits have also been detailed (Seifert et al., 2018a), leading to the identification of hybrid performance associated-sRNAs correlated with hybrid performance for grain yield (Seifert et al., 2018b). Thus, there is potential to incorporate sRNA expression into hybrid performance predictive models and achieve improved accuracy.

Despite these prior investigations, we have an incomplete picture of the genomic nature of sRNA variation. The relative variation among different genotypes or tissues for 21-nt, 22-nt or 24-nt sRNA abundance has not been investigated. In addition, we have limited understanding of the genomic features that contribute to consistent or variable sRNA abundance. While previous studies have examined sRNA profiles in maize and other plant species, they have often been limited to a single hybrid genotype or a single tissue. In this study, we combine sRNA profiles of five tissues from a diverse set of maize inbreds and hybrids to investigate genotypic and developmental variation in sRNA patterns, the levels and sources of sRNA variation, and the connection between sRNAs and heterosis.

## Results

Small RNAs were sequenced for five tissues (15 DAP endosperm, V7/8 leaf, V1 seedling root, V1 seedling shoot, and V7/V8 internode) of a panel of maize inbreds and hybrids (Supplemental Table 1). After filtering several samples with low sequencing depth, we obtained sRNA profiles for eight inbred and 13 hybrid genotypes for a total of 108 sRNA samples. An average of 5.2 million reads were generated per sample with a range of 1.9-17 million reads (Supplemental Table 2). These sRNA sequences were mapped to the B73 RefGen_v4 genome (Jiao et al., 2017) and then split by size class [bioinformatic approach detailed in (Stacey et al., 2016)]. Depending on the tissue, 15-34% of the sRNAs could be mapped uniquely to a single best location in the genome (Figure 1A). There are significant differences in the multi-mapping rates between tissues, for instance, nearly 50% of sRNAs map to highly repetitive sequences in endosperm and root compared to only around 30% in other tissues (Figure 1A, Figure S1A; ANOVA, Tukey’s HSD p < 0.001). When sRNAs map to multiple genomic locations the ‘count’ attributed to each location was scaled proportional to the number of locations to which it mapped.

**Figure 1.**
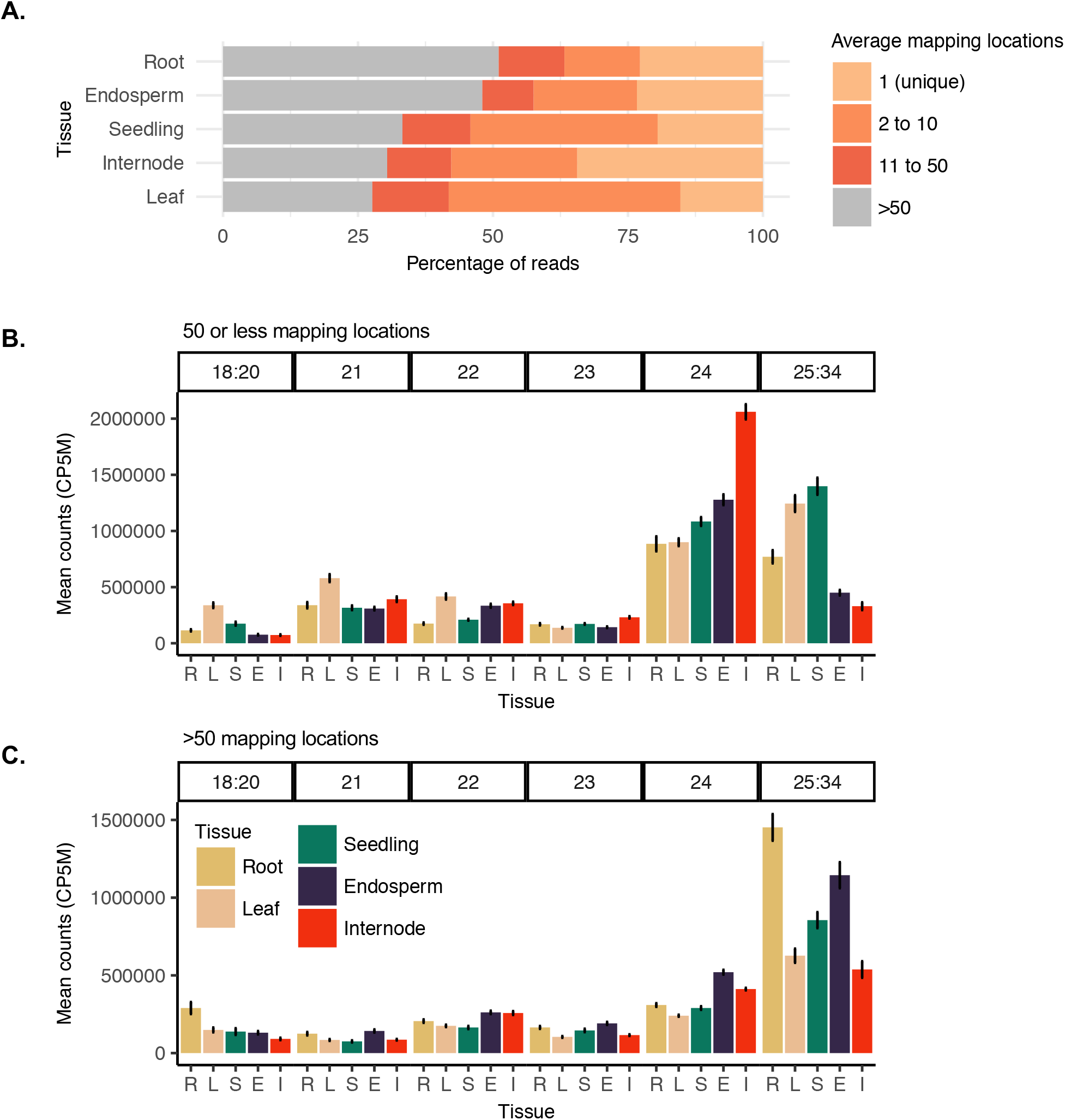
**A**.Average read multi-mapping frequency of small RNAs. Mapping frequency of genome mapped reads (excluding structural RNAs) of length 18 to 34-nt that perfectly aligned with no mismatches to the B73 reference genome for all samples in this study including inbreds and hybrids. For each sample mapping rates were categorised into four mapping frequency bins and expressed as a proportion of all mapped reads, then the distributions averaged for each tissue. Reads with mapping frequency 1 have a single (unique) high confidence mapping location, reads were then categorised into bins with 2 to 10, 11 to 50 or greater than 50 mapping locations. **B.** A comparison of the relative abundance of different size classes of small RNAs between different tissues for sequences with 50 or less mapping locations. **C** Relative abundance of sRNAs mapping to highly repetitive regions (>50 mapping locations). Error bars denote standard error among genotypes. R = “seedling-root” (n=20), L = “leaf” (n=18), S = “seedling-shoot” (n=21), E = “endosperm” (n=28), I = “internode” (n=21).

### Substantial developmental variation in sRNA profiles

The size distribution for sRNAs with 50 or fewer mapping locations was assessed for each tissue in the full set of inbred and hybrid genotypes (Figure 1B). Most work on sRNAs has focused on 21-nt, 22-nt or 24-nt size classes but there are also many sRNA reads that are 18-20-nt or 25-34-nt (Figure S2A). The abundance of individual size classes in the range 18-20-nt and 25-34-nt were low compared to 21-nt, 22-nt or 24-nt sRNAs (Figure S2A); however, collectively there were a substantial number of RNAs from these size classes when they are summed together (Figure 1B). There is significant variability in the total level of sRNAs between 25- and 34-nt observed in the five tissues. For some tissues (leaf and seedling shoot) these are the most abundant size class of sRNAs. If we normalize the sRNA expression levels based on the abundance of 25-34-nt sRNAs it is clear that 21-nt, 22-nt and 24-nt sRNAs are higher in internode than in other tissues (Figure S2B). Endosperm also tended to have somewhat elevated levels of 24-nt and 22-nt sRNAs compared to other tissues when normalised in this manner (Figure S2B). The analysis of sRNAs from highly repetitive regions (>50 mapped locations) revealed high contributions of 25-34-nt sRNAs and slightly elevated levels of 21-nt, 22-nt and 24-nt sRNAs in endosperm relative to other tissues (Figure 1C). We note that the choice about whether to include sRNAs from highly repetitive regions or sRNAs from the 25-34-nt size range in normalization can have significant impacts upon the perceived abundance of 21-nt, 22-nt and 24-nt, especially in comparisons among tissues. For this study, we elected to focus most analyses on sRNAs that have <50 mapped locations in B73 and to normalize using the total counts for the 18-34-nt set of sRNA size classes.

In order to evaluate variation in locus-specific abundances of sRNAs, we counted the 21-nt, 22-nt and 24-nt small RNA reads in 100 base pair (bp) windows of the maize genome that we termed “small RNA loci”. We examined in detail the genomic locations of expressed sRNA loci, which are defined as clusters with at least 10 counts per five million (CP5M) in at least one sample. sRNA loci can occur in both genic and non-genic regions of the genome. We divided the genome into five meta-feature categories by hierarchically classifying the genomic location of sRNA loci as non-coding RNA (e.g. miRNA and lncRNA), genic, gene-proximal (within 2 kb of a gene), transposable element (TE), or intergenic (greater than 2 kb from a gene). First, we determined the background distribution of each of the genomic features in the genome (Figure 2A). As expected, TEs accounted for the majority of the maize genome space (~66.7%) based on the latest TE annotation of B73 (Anderson et al., 2019), followed by intergenic regions (22.1%), genes and gene proximal regions (11.1%), and a very small fraction was annotated as non-coding RNA loci (0.1%). We then determined the distribution of genomic features represented by each size class of sRNAs (Figure 2B-D). For instance, for 21-nt sRNAs, relatively few non-coding RNA loci in the genome account for the majority of all 21-nt counts (Figure 2B). These analyses revealed substantial differences in the distribution of genomic features that contribute to the pool of 21-nt, 22-nt and 24-nt sRNAs that fit with expectations based on prior research on plant sRNAs. 21-nt sRNAs were mostly generated from non-coding RNAs, 22-nt predominantly from genes and TEs, while the largest fraction of 24-nt sRNAs comes from intergenic and gene proximal regions, with significant contributions from TEs as well.

**Figure 2.**
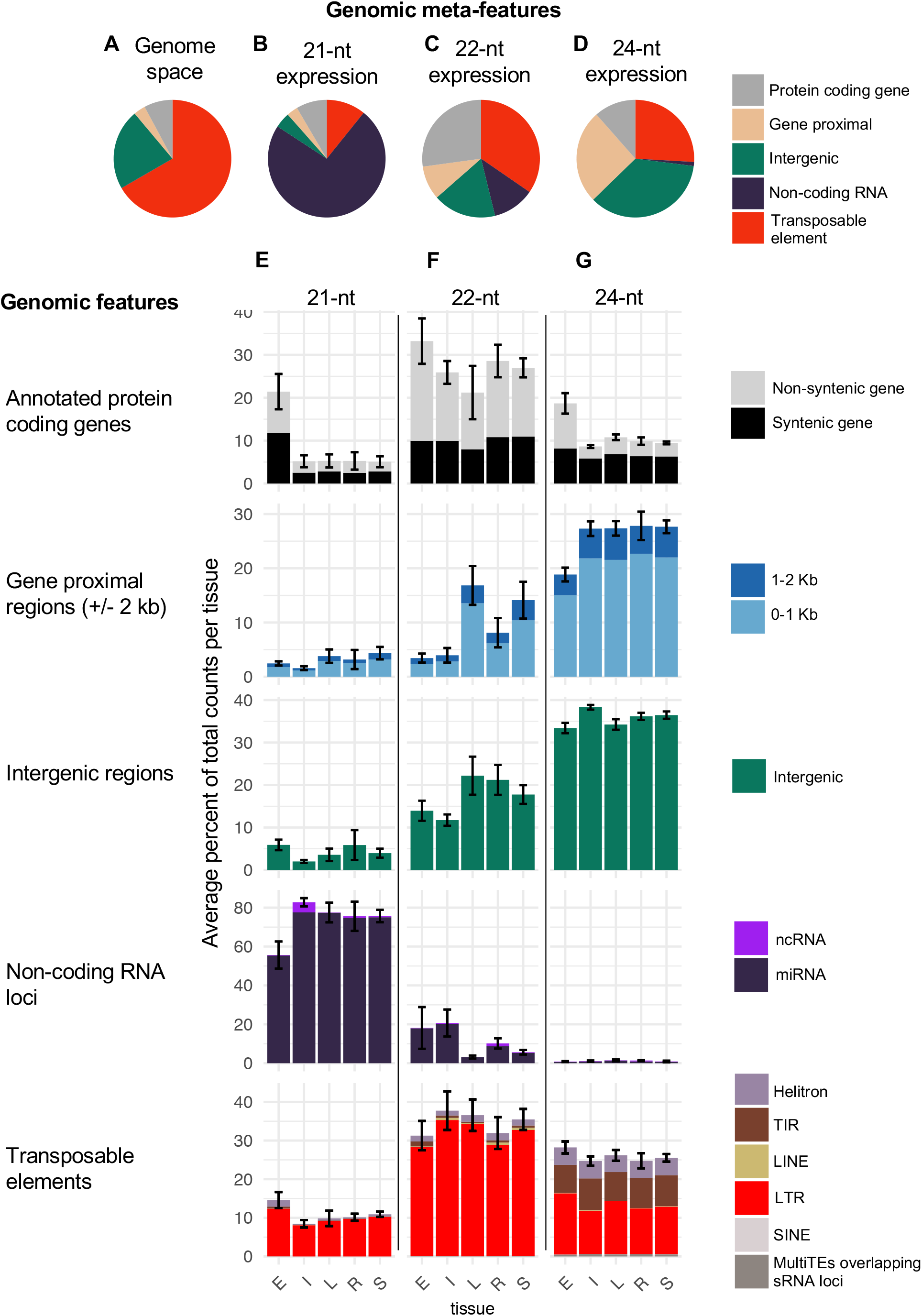
The distribution of sRNA expression among the genomic features of the genome. **A**. The background genomic distribution of genomic features was determined by annotating each 100 bp tile of the genome as either non-coding RNA (e.g. miRNA and lncRNA), genic, gene-proximal (within 2kb of a gene), transposable element (TE) or intergenic using the genome reference B73 RefGen_v4 and annotation based on Gramene version 36 and miRbase release 22. **B-D** The distribution of sRNA expression (CP5M) across meta-features for each sample was averaged to determine the meta-feature distribution for each size class. **E-G** Each meta-feature was then divided into constituent features and the average distribution of sRNA expression determined per tissue. The bars in the same vertical column add to 100%. Error bars denote SD among samples for the meta-feature category and represent the variation across the panel of inbred and hybrid genotypes, E = endosperm n = 28, I = internode n = 21, L = leaf n = 18, R = seedling-root n = 20, S = seedling-shoot n = 21. TIR = terminal inverted repeat; LINE = long interspersed nuclear element; LTR = long terminal repeat; SINE = short interspersed nuclear element.

It is unclear whether development leads to changes in the distribution of sRNA in the genome in maize. To explore the developmental variation in sRNA profiles, we assessed the relative contribution of distinct types of genomic regions to the sRNA expression profiles in each tissue (Figure 2E-G). For 21-nt sRNAs, vegetative tissues had a highly consistent profile, mostly comprised of miRNAs (Figure 2E). However, endosperm had a distinct increase in genic 21-nt abundance from both syntenic and non-syntenic genes and an increase in long terminal repeat (LTR, “RL”) retrotransposons. This was consistently observed for the different genotypes assessed in this study. For 22-nt expression, there was substantial variation between the tissues, particularly for gene proximal and intergenic regions (Figure 2F). Both endosperm and internode had substantially fewer 22-nt sRNAs in gene flanks, fewer in intergenic regions, but an increase in 22-nt miRNAs. In contrast, leaf and seedling-shoot had the highest level of gene proximal 22-nt sRNAs. This analysis also revealed that for the abundant class of genic 22-nt sRNAs, the majority are in non-syntenic genes (Figure 2F), which is in contrast to both 21-nt and 24-nt sRNAs (Figure 2E,G). For 24-nt sRNAs, again the vegetative tissues were relatively consistent and endosperm had a distinct increase in genic 24-nt counts. This analysis also highlighted that while 22-nt and 24-nt sRNA were both abundant at TEs, 22-nt were >90% LTRs while 24-nt were only around 50% LTRs with substantial numbers of reads mapping to both helitron and terminal inverted repeats (TIR) DNA transposons. Given that the amount of genome space occupied by LTR elements (59.5 %) is substantially larger than that of Helitrons (2.9 %) or TIR elements (3.0 %), there is a notable enrichment for Helitrons and TIR elements for 24-nt sRNAs.

### Globally, sRNA profiles are similar between inbreds and hybrids

The analyses above included both inbred and hybrid genotypes. Prior work in Arabidopsis (Groszmann et al., 2011) and maize (Barber et al., 2012) have found altered sRNA profiles in heterotic hybrids relative to their inbred parents suggesting this may be a general feature of heterosis (Greaves et al., 2015), although prior studies largely focused on a single hybrid genotype relative to the parents. We compared the profiles of the sRNAs in each of the tissues from the full set of inbred and hybrid genotypes used in this study (Figure 3A). Using a two-way ANOVA interaction model for unbalanced design (Type-III sum of squares), we found that pedigree (inbred/hybrid) is not a significant factor in sRNA counts for any mapping rate category (ANOVA, pedigree p > 0.05). Likewise, there was limited evidence for any significant global difference in the abundance of the different sRNA size classes specifically between inbreds and hybrids, for example in seedling-shoot tissue (Figure 3B) or other tissues (Figure S3). In addition, the distribution of the types of genomic loci contributing to sRNA expression was nearly identical between inbreds and hybrids for all size classes (Figure S4). These analyses suggest that maize hybrids exhibiting substantial heterosis do not have globally unique sRNA compositions relative to the inbred parents.

**Figure 3.**
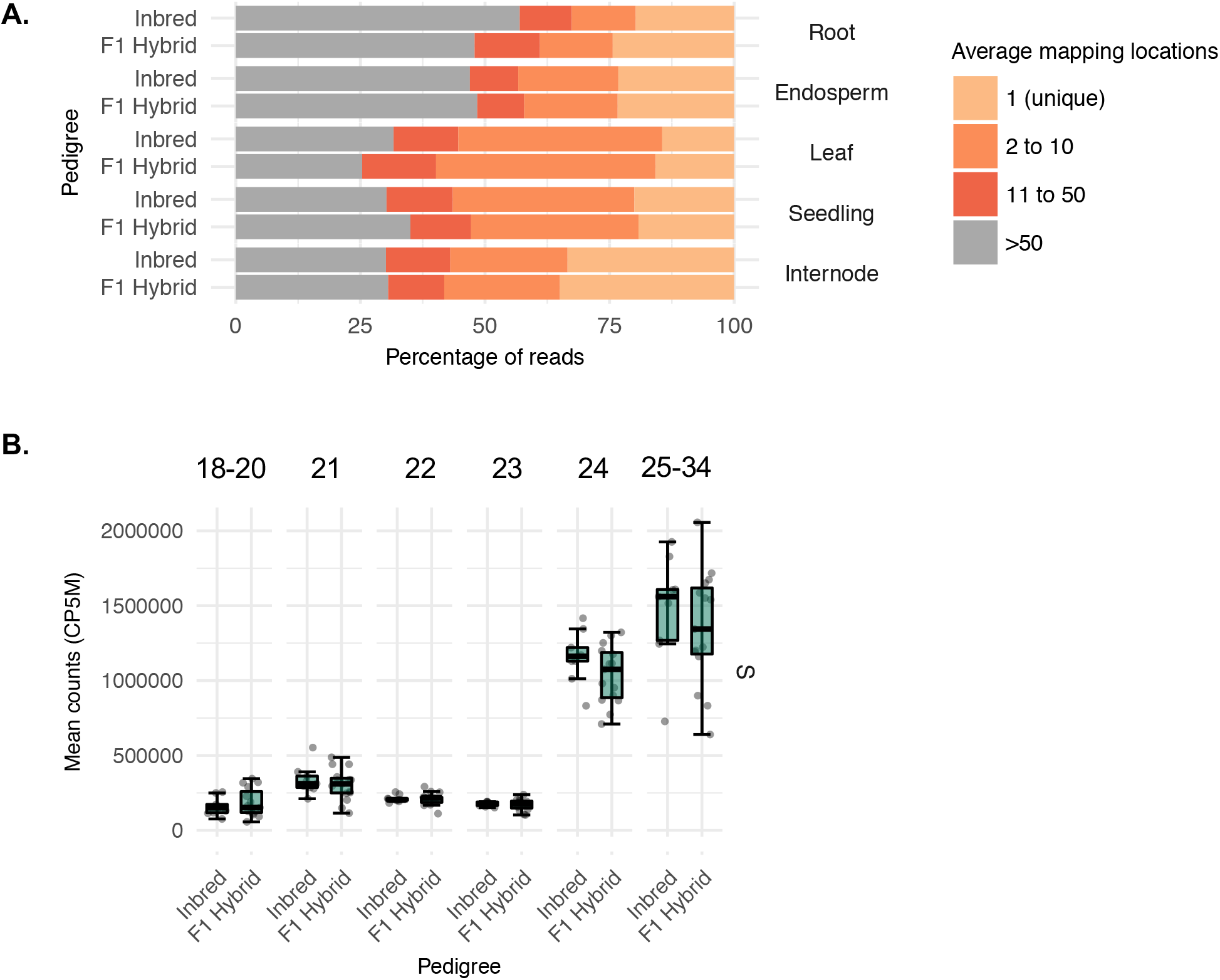
**A.** Comparisons of multi-mapping rates in inbreds relative to hybrids. The average read multi-mapping frequency for all genome-matched small RNAs was summarised into four categories and then divided into averages for inbreds and F1 hybrids. Bars represent the average of all genotypes for each sample. **B.** A comparison of sRNA abundance between inbreds and hybrids for seedling-shoot tissue. Each dot represents a genotype, whiskers correspond to the first and third quartiles (the 25th and 75th percentiles). S = “seedling-shoot” (n = 21).

### Size class-specific developmental and genetic variation in sRNA expression

Prior work has suggested that sRNA profiles may capture unique information missed by SNPs or gene (mRNA) expression data (Seifert et al., 2018a, 2018b). For each hybrid and inbred sample, gene expression levels were profiled by RNA-seq using the same RNA sample used for small RNA analysis (Li et al., 2019). To compare and contrast sRNA profiles with gene expression we summarised RNA-seq reads to counts per million (CPM) per B73v4 gene loci (APGv4 release 36). The mRNA and sRNA levels per loci were used to perform principal component analysis (PCA) to assess the relationships of profiles for different tissues and genotypes (Figure 4). For the mRNA data, variation in the expression of genes was heavily driven by tissue type (Figure 4A), as has been observed previously [e.g. (Zhou et al., 2019)]. Samples clustered by tissue-type into discrete groups with PC1 explaining 28% of the variation and PC2 another 17%. For 21-nt and 22-nt sRNA loci, samples also clustered by tissue (Figure 4B-C). However, seedling-shoot and leaf were not separated by the first two PCs, which also accounted for relatively less of the variation compared to mRNA data capturing ~16% and 9% respectively. In contrast to mRNAs and 21-22-nt sRNAs, 24-nt sRNA loci were not strongly clustered by tissue type and instead there was evidence for separation by genotype, particularly in PC2. Separate PCAs were performed for each type of molecule (mRNA, 21 nt, 22 nt or 24 nt) within each tissue (see representative examples in Figure 4E-F, Figure S5). One hypothesis is that heterosis could lead to a consistent change in sRNA profiles in hybrids relative to their parents. However, there was no evidence for clustering of sRNA profiles of hybrids relative to inbreds, instead in most cases the hybrids are intermediate relative to the two parents in both PC1 and PC2; an example is highlighted using dashed lines in Figure 4 F for the PH207 hybrids. Very similar patterns are seen for the other size classes and tissues (representative examples provided in Figure S5). This behaviour of the hybrids (intermediate to the parents) was less clear for gene expression (Figure 4E).

**Figure 4.**
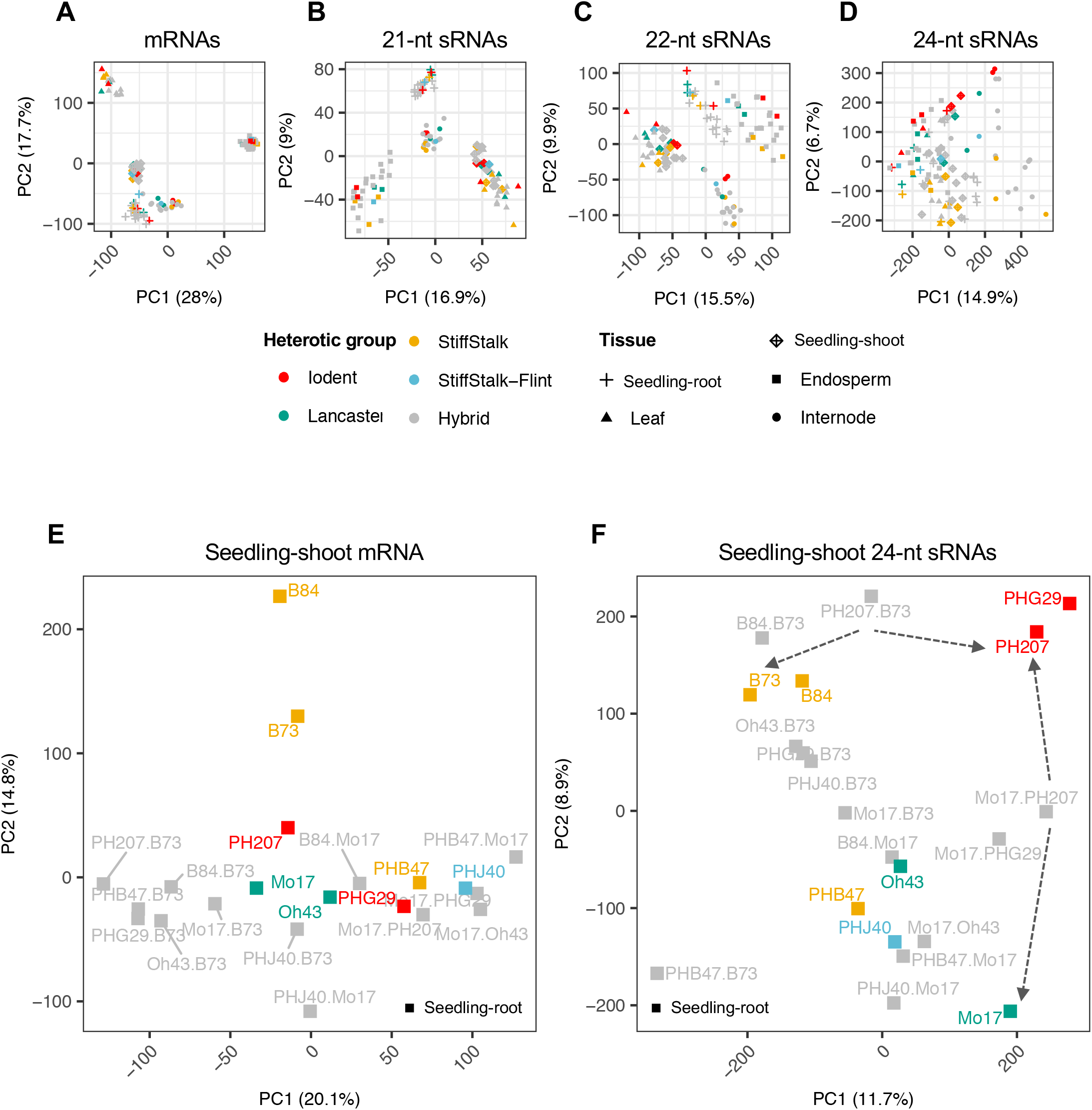
**A-D.** Principle component analysis (PCA) of all tissues on expressed loci for mRNA and 21-nt to 24nt sRNAs. **E.** PCA specifically for seedling-shoot mRNAs, and **F.** seedling-shoot 24-nt siRNA. For each sRNA size class PCA was performed using expressed loci with greater than 10 CP5M in at least one sample; while mRNA loci with greater than 1 CPM were used as input. Counts were log2 transformed, scaled by unit variance and clustered using singular value decomposition. Colours represent genotype (heterotic group or hybrid), symbols represent tissue type. Percentages in brackets refer to the percent of variance explained by each PC. The dashed lines provide an example relationship between two PH207 hybrids and their respective parental inbreds.

### Inbred-specific sRNA expression drives sRNA variability among genotypes

Next, we examined sRNA loci for variable expression between inbred parents. The experimental design did not include biological replicates, limiting our ability to robustly assess differential expression. However, we compared trends in terms of the proportion of sRNA loci with >2-fold and >4-fold variation or Single Parent Expression (SPE) among inbreds, to determine whether certain size classes or tissues might have more variation (Figure 5A-D). To stringently assess variable expression, only loci with greater than 10 CP5M (or the equivalent 2 CPM for mRNA) in at least one genotype were considered and SPE was strictly defined as greater than 10 CP5M in one parent and 0 in the other. The proportion of sRNA loci with variation for sRNA expression (Figure 5B-D) is much greater than the proportion of genes that exhibit variation (Figure 5A). Across all tissues and sRNA size classes, ~50% of regions with sRNA expression had at least 4-fold variation and approximately half of these were SPE in at least one pair of inbreds. In total, >60% of all sRNA loci had >2-fold variation; by contrast many fewer genes exhibited variation (Figure 5A). The level of variation was slightly higher for internode than for other tissues for 21-nt and 22-nt sRNAs, but was slightly lower in this tissue for 24-nt sRNAs. For leaf, root and seedling-shoot tissue the 24-nt sRNAs seem to have more variable loci than for 21-nt or 22-nt sRNAs while the relative levels are similar for the size classes for internode and endosperm tissue.

**Figure 5.**
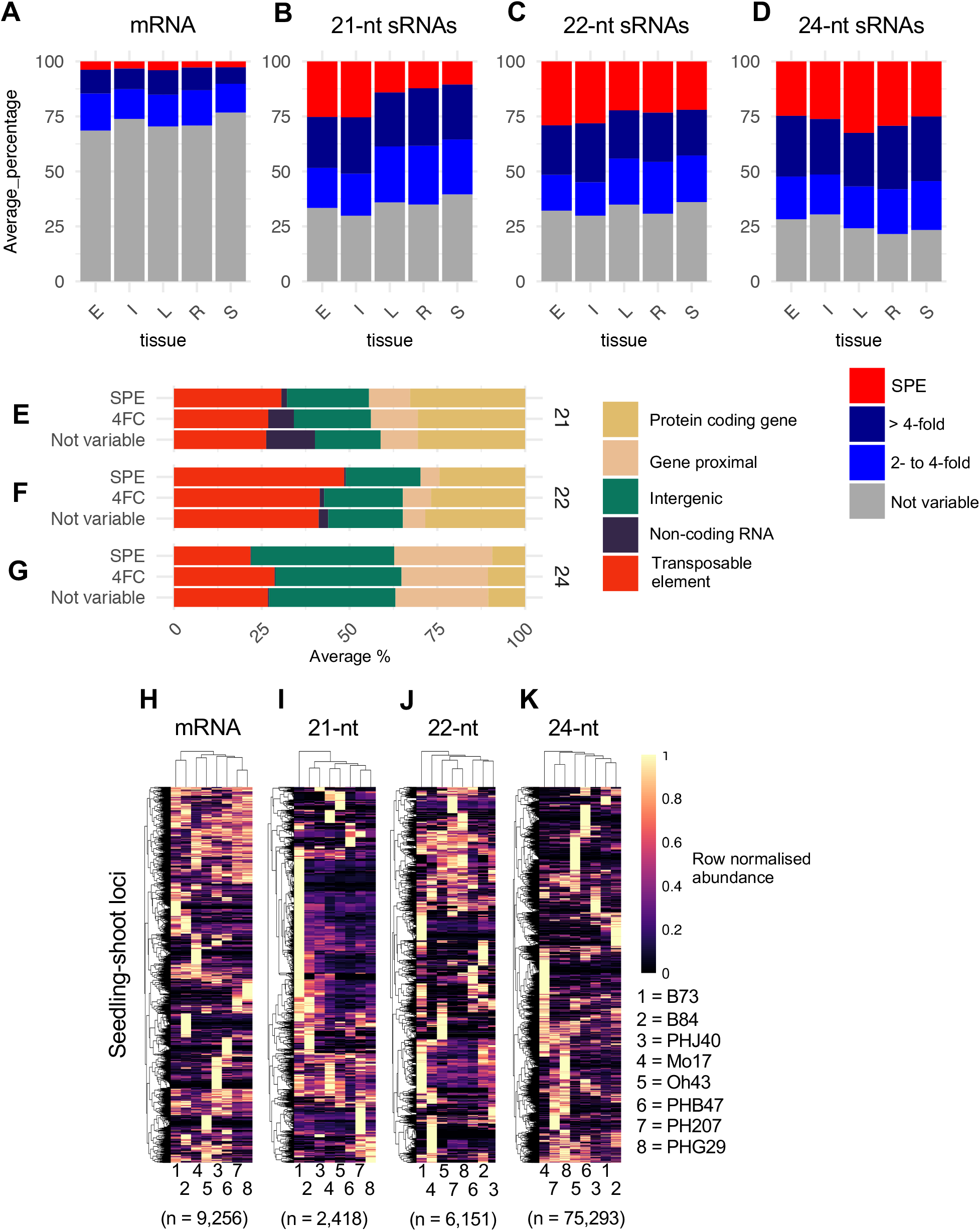
**A-D.** Frequency and source of expression variation between inbreds. For each hybrid the variability in expression between the parents was assessed. Only loci expressed in at least one of the parents were considered (10 CP5M or greater in one parent) by calculating the log2 ratio of high parent/low parent. The proportions of loci in each variable expression category (“not variable”, “2-4 fold”, “> 4 fold” and “SPE”) were collated and then averaged per tissue and size class combination. SPE strictly defined as > 10 CP5M in one parent and 0 in the other parent. R = “seedling-root” (n=13 contrasts), L = “leaf” (n=11 contrasts), S = “seedling-shoot” (n=13 contrasts), E = “endosperm” (n=20 contrasts), I = “internode” (n=13 contrasts). **E-G** Genomic loci contributing variable expression of sRNAs. The distribution of sRNA expression (CP5M) across meta-features for each sample was averaged per tissue and per variable expression category to determine the meta-feature distribution for each size class. n = 70 contrasts averaged per size class. **H-K** Expression patterns of loci with variable expression in seedling-shoot. Per size class, each loci with variable expression (>4-fold or SPE) in at least one contrast was profiled for each hybrid. Expression was normalised to the maximum expression (0-1 scale) for that loci and hierarchically clustered. The number of variable loci per size class is show in parenthesise, for visualisation a maximum of 40,000 were included per heatmap.

Considering the loci with variable sRNA expression, we hypothesised that certain categories would be more likely to vary between inbreds; for instance, sRNAs in genic regions might increase or decrease in concert with changes in gene expression, while TE-associated sRNAs might be highly consistent between genotypes, given that TEs generally remain repressed in the inbreds profiled. However, an analysis of the genomic loci contributing variable sRNAs (Figure 5E-G) revealed that TEs, gene loci and intergenic regions were, broadly speaking, equally as likely to be variably expressed. For 21-nt loci, the relatively few non-coding RNA loci - mostly miRNAs - were rarely SPE and relatively less likely to be > 4-fold variable, which led to increases in the proportion of the other genomic feature categories (Figure 5E; S6A). The 22-nt loci showed slightly more variation between variable and not variable loci, particularly in genes and TEs (Figure 5F); this was most evident when comparing tissues (Figure S6B). For instance, in endosperm and internode, variable (>4-fold and SPE) expression patterns are relatively enriched in TEs and depleted in genes (Figure S6B). For 24-nt sRNAs there was very little difference between the proportion of genomic features of variable (>4-fold and SPE) compared to not variable loci consistently across tissues and the individual inbred comparisons (Figure 5G, S6).

For each size class and tissue, we identified a set of non-redundant loci that exhibit at least 4-fold variation, including SPE, in at least one pairwise comparison. For any given loci we assessed variation across multiple inbred contrasts both within and between heterotic groups (Figure 5H-K, S7). Variable expression could reflect consistent differences between heterotic groups, rare gains, or rare losses of expression. A heatmap of the normalized expression levels for seedling-shoot loci revealed that many sRNA and mRNA loci have expression in only one of the inbreds (Figure 5H-K) such that expression variation was driven by rare gains of expression. While rare gains in expression were the most common type of variation, there were differences in the number of loci, or genes, that exhibited expression in the majority of inbreds. For mRNA and 22-nt sRNAs there was a set of genes or loci that exhibited expression in the majority of genotypes but this pattern was quite rare in 21-nt and 24-nt sRNAs. There are also differences in the level of bias towards the B73 reference genome. There are a large number of 21-nt sRNAs that are expressed only, or primarily, in B73. In contrast, there was very little enrichment for B73-specific expression for genes or 24-nt sRNAs. These overall patterns of variation among genotypes were quite similar in other tissues for both mRNA and 21-24-nt sRNAs (Figure S7). Thus, we found that the majority of sRNA loci showed variation in expression; and this was predominantly due to high levels of expression in a single inbred.

There was also a high level of expression variation for sRNAs among genotypes within a single tissue. We were interested in assessing how the variation observed in a single tissue related to expression levels and patterns in other tissues (Figure S8). For each of the heatmaps of variation among inbreds (shown in Figure 5H-K), we maintained the same order of genes and genotypes and assessed the expression of the same loci in the other tissues (Figure S8). The analysis of 21-nt sRNAs with variable expression in seedling-shoot tissue revealed that very few of these loci exhibit similar genotype variation in other tissues. Many of these were expressed at substantially lower levels or exhibited fairly consistent expression among genotypes in other tissues (Figure S8A). For instance, the block of B73-specific loci marked as cluster A in Figure S8A is rarely expressed in other inbreds or in B73 in other tissues (with the exception that these loci are lowly expressed in leaf tissue which is the most similar tissue type to seedling shoot). By contrast, for both 22-nt and 24-nt sRNA loci, the patterns of variability observed in seedling-shoot were frequently seen in other tissues, albeit at slightly lower levels of expression (Figure S8B-C).

### sRNA loci exhibit non-additive expression across a panel of different hybrids

We sought to examine if and how hybrids differ from inbreds for sRNA expression. We focused on two separate types of hybrid-inbred comparisons, with comparisons conducted discretely for each inbred parent triplet in this study then summarised across triplet comparisons. An initial analysis focused on searching for loci that were only expressed in the parents or in the hybrid. The majority of loci that were expressed in parents were also expressed in hybrids but there was a small subset of less than 1% of loci that were only expressed in the inbred parents or only in the F_1_ hybrid (Figure 6A-D). Hybrid-specific expression was more common than inbred-specific expression for both genes and sRNA loci; however, hybrid-specific expression was an order of magnitude more rare for gene expression compared to sRNA expression (Figure 6A). sRNA loci expressed only in the hybrid or the parents were located in a variety of genomic features including TEs, genes and intergenic regions (Figure 6E). There were slightly higher levels of inbred-specific or hybrid-specific expression for 24-nt sRNAs compared to other size classes (Figure 6B-D).

**Figure 6.**
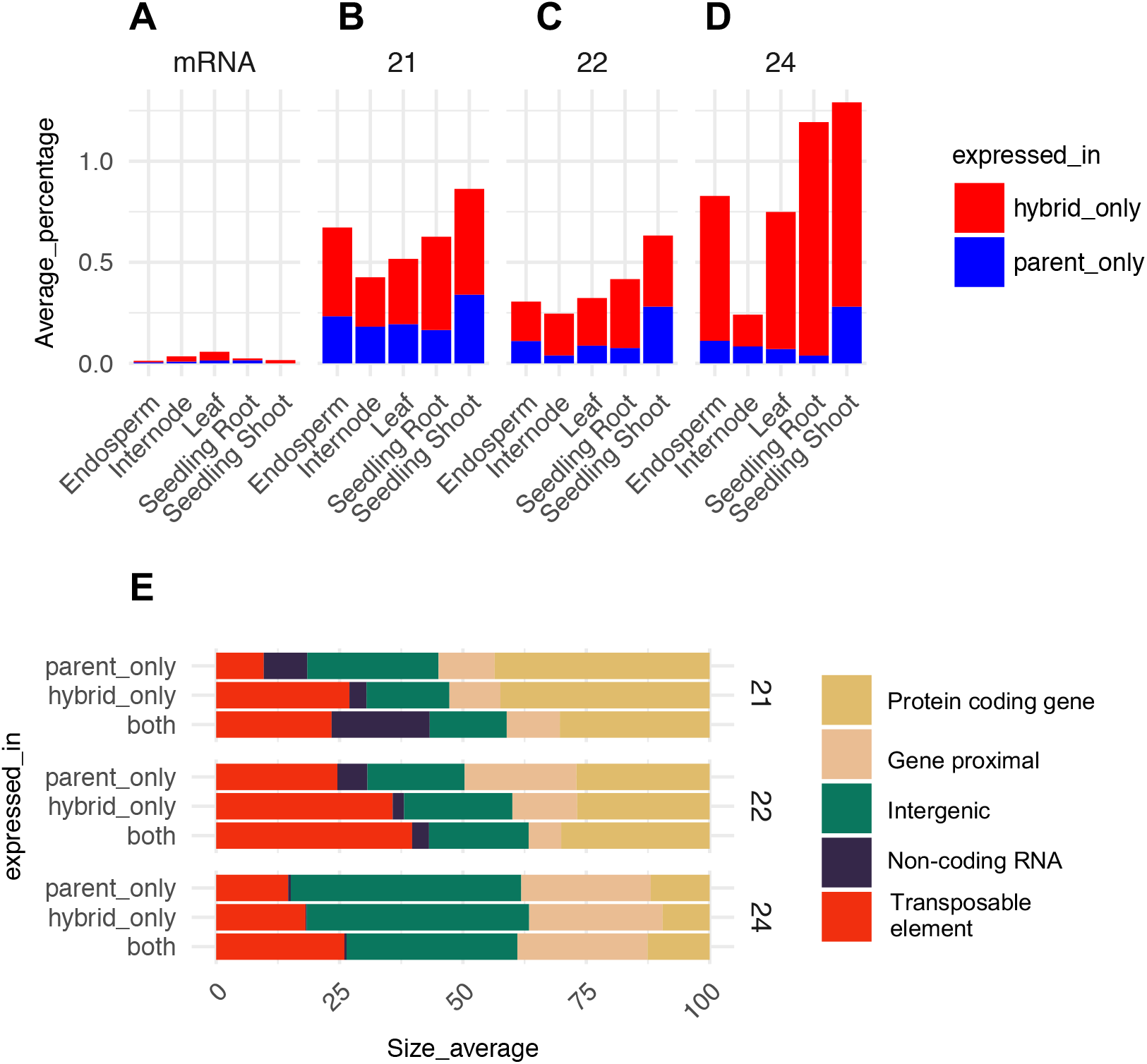
**A**. Proportion of loci uniquely expressed in hybrids. For each inbred hybrid trio the proportion of loci expressed only in either hybrids or inbred parents was calculated and then proportions averaged for all sample groups in each tissue. For sRNAs only loci expressed >= 10 CP5M in one of each trio were considered, “hybrid only” defined as a loci >= 10 CP5M in the hybrid and < 2 CP5M in both parents; “parent only” defined as >= 10 CP5M in at least one parent and < 2 CP5M in the hybrid. **E.** The distribution of sRNA expression (CP5M) across meta-features for each sample was averaged per tissue and per expression category to determine the meta-feature distribution for each size class. N = 38 contrasts averaged per size class.

For loci expressed in both inbreds and hybrids, we compared the hybrid expression levels to the average of the two parents (the mid-parent level; Figure 7A). This analysis revealed a normal distribution of hybrid expression values relative to mid-parent levels, with examples of novel increases or reductions of expression in the hybrids. Overall, for sRNA loci that did not vary between parents, the levels of sRNA abundance exhibited a fairly normal distribution centered at additive levels, similar to the distribution for mRNA although mRNA values had less variance. This trend was consistent across all tissues and size classes (Figure S9A); although, in root 21-nt loci were shifted towards below parental levels, while 22-nt and 24-nt loci were shifted above parental levels. We proceeded to address the additivity of expression levels for sRNA loci with differential abundance in the inbred parents. Prior work on transcript abundance suggests that most genes that are differentially expressed in the two parents exhibit additive, mid-parent (MP), patterns in the hybrids with some examples of expression at the high-parent (HP) or low-parent (LP) levels and we find similar results for mRNA data for the parents and hybrids in this study (Figure 7B-D, dashed line). Likewise, sRNA loci exhibit a similar frequency of non-additive expression in all categories: 2-to 4-fold variable, greater than 4-fold and SPE loci (Figure 7B-D, solid lines). In several tissues, the 21-nt sRNAs exhibited some enrichment for expression values that were lower than the mid-parent values. While the relative frequency of non-additive expression was similar for gene expression and sRNA expression, given that many more sRNA loci were variably expressed between the parents, a much higher proportion of sRNA loci overall exhibited non-additive expression.

**Figure 7.**
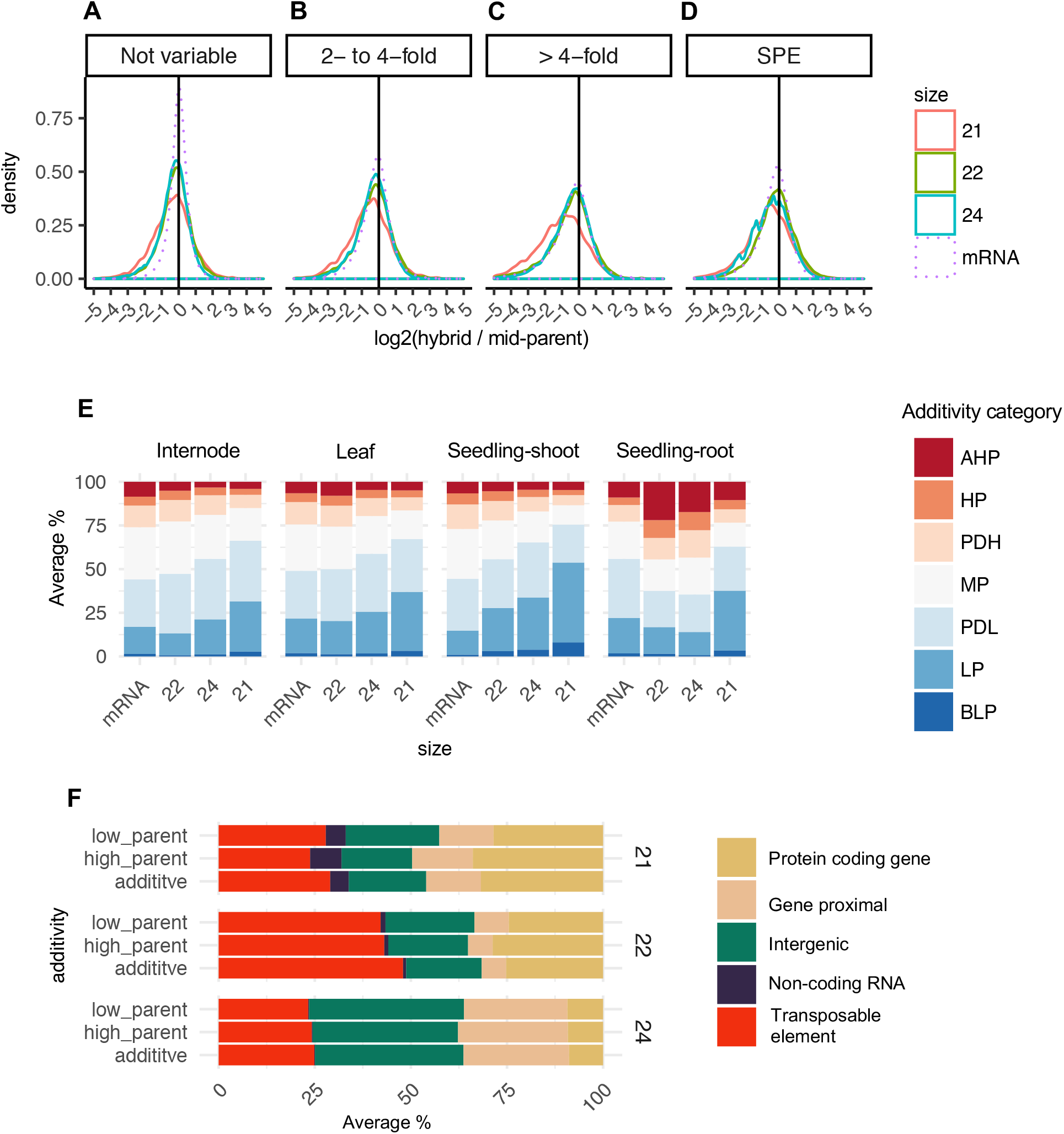
**A-D.** Overall distribution of additivity in hybrids per sRNA size class compared to mRNA. Additivity was calculated as log2(hybrid / mid-parent) for sRNA and mRNA loci with at least 10 CP5M (2 CPM for mRNA) in one parent and divided into categories for the level of expression variation between the inbred parents. SPE = single parent expression. For plotting the tails were concatenated at +/− 5. **E.** Distribution of non-additivity categories per tissue (d/a values). For sRNA loci that are 4-fold or greater variable between inbred parents the degree of dominance (d/a) calculated as [hybrid – mid-parent/(high-parent – low-parent/2)] was determined. d/a values were then divided into either additive mid-parent or 6 non-additivity levels: Above High Parent (AHP) > 1.25; High Parent (HP) 0.75: 1.25; Partial Dominance High Parent (PDH) 0.25: 0.75; Mid-parent (MP) −0.25: 0.25; Partial Dominance Low Parent (PDL) −0.75: 0.25; Low Parent (LP) −0.75: −1.25; Below Low Parent (BLP) < −1.25. **F** The genomic feature of non-additive loci. For non-additive high-parent like (AHP, HP and PHD) and low-parent like (PDL, LP, BLP), the distribution of sRNA expression (CP5M) across meta-features for each sample was averaged per tissue for each size class. N = 30 hybrid parent trio contrasts averaged per size class, Endosperm was excluded owing to the genome imbalance.

Prior analyses have found an enrichment for low parent expression for sRNAs (Groszmann et al., 2011; Shen et al., 2012; Barber et al., 2012). In this dataset, for both sRNAs and genes, a greater number of loci were below mid-parent levels in the hybrids (Figure 7E, blue bars). However, for internode, leaf and seedling-shoot tissue, sRNA loci showed a greater level of low-parent expression compared to mRNA, consistent across 21-to 24-nt size classes. Similar trends were observed in endosperm after taking into consideration the genome imbalance in this tissue (Figure S9B). By contrast, 22-nt and 24-nt loci in root tissue had higher levels of high-parent expression compared to other tissues and compared to gene expression in root tissue. Next, we examined the genomic location of non-additively inherited sRNA loci. This analysis revealed that non-additive 21-nt, 22-nt and 24-nt loci occur in all regions of the genome (Figure 7F), largely reflecting the expression distribution of these sRNAs (Figure 2; Figure 5 E-G). The patterns were highly consistent between high-parent-like and low-parent-like loci. While there was substantial variation between the size classes, overall, there was very little difference in the distribution of genomic features between additive and non-additive loci (Figure 7F), similarly the different tissues exhibited very consistent genomic feature distributions (Fig S10), with the exception that 22-nt loci in internodes were somewhat enriched for TEs compared to other tissues. Overall, these results suggest that sRNA loci originating from different genomic features have a similar propensity to be non-additively expressed.

## Discussion

sRNAs have been hypothesized to play key roles in plant responses to environmental conditions, transgressive inheritance and heterosis (Greaves et al., 2015; Ryder et al., 2014; Li et al., 2017). However, deciphering their role in crops such as maize has been limited by our incomplete picture of the sources of sRNA variation. Many studies have focused on a single tissue and/or a limited number of genotypes. We were motivated to undertake a broad survey of sRNA profiles in several distinct tissue types in a panel of diverse inbreds and hybrids in the highly heterotic crop, maize. We performed a detailed analysis of sRNA and mRNA expression in a panel of 108 maize samples including 8 inbreds and 14 hybrids across five tissues: endosperm, seedling root, leaf, seedling shoot and internode. This has provided a more detailed view of the genomic features that contribute sRNAs of different size classes, the developmental variation for sRNA abundance and the potential connections between sRNAs and heterosis.

### Genomic features that contribute to consistent or variable sRNAs

We documented substantial developmental and genotypic influences on sRNA variation; however, we found that the contributions of these factors were different for 21-nt, 22-nt and 24-nt sRNA expression. This likely reflects established differences in molecular functions and modes of biogenesis of each size class. For instance, 21-nt sRNAs are predominantly miRNAs originating from a relatively small number of non-coding miRNA loci, 24-nt clusters are frequently located near genes, and 22-nt clusters often originate from repetitive sequences (Wei et al., 2009; Nobuta et al., 2008; Regulski et al., 2013; Barber et al., 2012). While these generalizations are supported by our analyses, we also find that each size class of sRNAs has a complex make-up that can vary in different tissue types. For example, while 22-nt sRNAs are enriched for TEs (~30-40% of 22-nt sRNA loci) they also are frequently associated with genes, especially non-syntenic genes (Figure 2). 21-nt sRNAs are rarely found associated with genes, except in endosoperm tissue where ~20% of 21-nt sRNAs arise from genes. Overall, there was more variability in the features associated with 22-nt sRNAs in different tissues compared to 21-nt or 24-nt sRNAs (Figure 2E-G). In particular, we note the reduction in 22-nt sRNAs in gene flanks in endosperm and internode and also that the abundant class of genic 22-nt sRNAs mostly occur in non-syntenic genes. One possibility is that many of these non-syntenic genes could be nonfunctional and silenced by the plant; the enigmatic class of 22-nt siRNAs in maize might be involved in this silencing mechanism. Previously, Barber et. al (2012) observed differences in sRNA abundance between retrotransposon families. Here, armed with an updated annotation of maize TEs (Stitzer et al., 2019; Anderson et al., 2019), we observed differences in the distribution of 22-nt and 24-nt sRNA between different TE orders with 22-nt around >90% LTRs while 24-nt sRNAs have a high composition from helitron and TIR DNA transposons, far greater than the genome space these TEs occupy; 2.9% and 3.0% respectively. This hints at differences in the potential roles of sRNAs in regulating different classes of TEs in maize.

We also performed comparisons of multiple inbreds or of inbreds with hybrids to identify sRNAs that exhibit variable expression between different genotypes or non-additive expression in hybrids relative to parents. While there were many examples of sRNA loci that exhibited variability in expression levels, we did not find that particular types of sRNAs were more likely to exhibit variation. The distribution of genomic features associated with variable 21-nt, 22-nt or 24-nt sRNAs were largely similar to the complex distribution observed for sRNA loci without variable expression. This suggests that specific types of sRNAs are not necessarily enriched for variability.

### Varying influence of tissue and genotype on different size classes of sRNAs

Prior work on sRNAs and heterosis has often been limited to a single hybrid genotype or a single tissue, for instance in maize (Barber et al., 2012; He et al., 2013; Regulski et al., 2013; Xin et al., 2014) and Arabidopsis (Groszmann et al., 2011; Shen et al., 2012; Li et al., 2012, 2015b). A significant motivation of this study was to investigate whether prior interpretations are representative of other tissues and developmental stages or whether there is variation and uniqueness between tissue types. The analysis of variability for sRNA expression in different tissues and genotypes revealed distinct patterns for different size classes of sRNAs. Gene (mRNA) expression has greater variability among tissues than among genotypes. For example, gene expression profiles for the B73 root were more similar to the root samples of other genotypes than to profiles from B73 leaf; this trend was generally consistent across genotypes and tissues. A very similar pattern was also observed for 21-nt sRNAs and to a lesser extent 22-nt sRNAs (FIgure 4). This suggests substantial differences in tissue-specific abundance of 21-nt and 22-nt sRNAs that are largely reproducible in different genotypes. In contrast, 24-nt sRNAs exhibit relatively little clustering by tissue type and instead are clustered by genotype. This indicates a greater role for genetic variation in driving differences in 24-nt sRNA abundance. This is supported by the analysis of the patterns of variable abundances for the expression of sRNA loci in multiple tissues. 21-nt sRNAs that exhibit variable expression seem to often have tissue-specific expression while 24-nt sRNA variation is more frequently observed in multiple tissues (Figure S8). Thus, differences in 24-nt sRNA abundance in one tissue are more predictive of variability in other tissues while observations of 21-nt or 22-nt sRNAs will often be limited to a single tissue type. Thus, developmental variation is a significant factor that should be considered when designing or interpreting analysis of heterosis and sRNA profiles.

### Connections between sRNAs and heterosis

Important early work on the inheritance of sRNA in hybrids in Arabidopsis made the observation that F_1_ hybrids have reduced levels of 24-nt siRNAs relative to parents for many loci, with an estimated 25% or greater drop in production of these siRNAs (Groszmann et al., 2011). Similarly, a study in maize suggested that 24-nt sRNAs often exhibit non-additive expression in hybrids with expression levels that were lower than expected (Barber et al., 2012). Here, we find that on a global scale over a diverse panel of heterotic hybrids in maize there is little difference between the global profiles of 21-nt, 22-nt or 24-nt sRNAs between inbreds and hybrids (Figure 2B, Figure S3). Likewise, clustering of sRNA profiles did not reveal any underlying characteristic of hybrids that could separate them as a class from inbred parents (Figure 4). This is consistent with the data underlying other work, noting that overall there appears to be no general shift in the abundance of the size classes between inbreds and hybrids in maize (Barber et al., 2012) or Arabidopsis (Li et al., 2012). There are many sRNAs with non-additive expression levels but the majority of these are expressed at levels expected for partial or complete dominance with rare expression outside of the parental range. There was no strong enrichment for sRNA loci with non-additive expression outside the parental range that was consistently observed in multiple tissues (Figure 7E). Collectively, our results and this body of literature suggest that hybridization likely does not substantially alter biogenesis or accumulation of different sRNA size classes.

In addition to comparing the global profiles of 21-nt, 22-nt or 24-nt sRNAs in inbreds and hybrids, we also performed targeted searches for sRNAs that exhibit unexpected abundance in hybrids relative to parents. There was an enrichment for sRNAs solely expressed in hybrids or inbreds relative to mRNAs, but these still accounted for only ~1% of sRNA loci (Figure 6). In addition, these sRNA loci with unique hybrid expression patterns did not necessarily show strong enrichments for certain genomics features. There was no evidence that genic or TE-derived sRNAs experience unusual accumulation patterns within hybrids relative to parents. Similarly, the analysis of non-additive expression for sRNAs in hybrids did not find striking enrichments for specific features. These findings suggest that sRNA are unlikely to be the general basis of heterosis. Clearly, quantitative variation in sRNA expression is common and the non-additive expression of particular loci could be important for heterotic traits. However, we did not find evidence of whole-scale alterations to sRNA accumulation or hybrid uniqueness that would suggest a major upheaval of sRNA regulation and influence in hybrids.

Several prior studies on maize have provided interesting insights into heterosis and sRNA abundance. Barber et al. (2012) reported non-additive expression, particularly for 24-nt sRNAs from repetitive genome regions, in maize hybrids, but also found that mutations that greatly reduced the total abundance of 24-nt sRNAs did not greatly reduce heterosis. Seifert et al. (2018a, 2018b) assessed variation in seedling sRNAs in a panel of 21 inbred lines and found many sRNAs (often 22-nt or 24-nt sRNAs) that were associated with grain-yield heterosis in hybrid lines. We found quite high levels of variation for 22-nt and 24-nt sRNAs among genotypes. Given the large number of sRNA loci and the high proportion of variation this means that there are many more variable sRNA loci than differentially expressed genes. For a biomarker with high levels of variation it is not surprising that a subset of variation patterns would match variation for a measured trait. It is unclear whether this association was due to a causal relationship between sRNA abundance and yield heterosis or whether these observations are simply a reflection of the high level of variation.

This study provides a detailed understanding of the genomic regions that are associated with sRNAs in maize. We confirm substantially different profiles of different size classes of sRNAs in distinct tissues of maize and find differences in the genomic regions that contribute these sRNAs in distinct tissues. Detailed comparisons of inbred and hybrid sRNA profiles helps to provide a detailed understanding of the potential roles of sRNAs in heterosis. We fail to find evidence for major shifts in sRNA abundance that would provide clear insights into the core mechanisms of heterosis.

## Materials and Methods

### Plant material

To represent the maize heterotic groups, inbred lines were selected from the stiff stalk synthetic group (B73, B84, PHB47, PHJ40), the non-stiff stalk synthetic group (Mo17, Oh43) and the iodent group (PH207, PHG29). This investigation was also part of a larger germplasm sampled for mRNA analysis (Li et al., 2019). Hybrids were generated by crossing each of these selected inbred lines by three male genotypes that included B73 (stiff stalk synthetic), Mo17 (non-stiff stalk synthetic), and PH207 (iodent) in the scheme described in Supplemental Table 1. Five tissues were sampled from the inbred and hybrid genotypes including seedling root at Vegetative 1 (V1) stage (“Seedling-root”, “R”), seedling shoot at V1 (“Seedling-shoot”, “S”), the middle of the eighth leaf at V7/8 (“Leaf”, “L”), the upper most elongated internode at V7/V8 (“Internode”, “I”), and endosperm at 15 days after pollination (“Endosperm”, “E”) (Li et al., 2019). Seeds were planted at the Minnesota Agricultural Experiment Station located in Saint Paul, MN on 05/16/14 with 30 inch row spacing at ~52,000 plants per hectare and sampled during the 2014 field season. For the V1 tissues (Root and Shoot samples), seeds were planted in Metro-Mix300 (Sun Gro Horticulture) with no additional fertilizer and grown under greenhouse conditions (27C/24C day/night and 16 h light/8 h dark) at the University of Minnesota Plant Growth Facilities during 2014.

### sRNA-seq library construction, sequencing and analysis

Samples were flash frozen in liquid nitrogen and the small RNA enriched total RNA fraction was extracted using the miRNAeasy Mini Kit (Qiagen); this preparation was split and was used for both sRNA-seq and RNA-seq. Extracted RNA was DNase treated using the TURBO DNA-free kit (Life Technologies). Sequence libraries were prepared by the Joint Genome Institute following standard the TruSeq Small RNA library preparation protocol (Illumina, San Diego, CA). Samples were sequenced on an Illumina HiSeq 2500 (Illumina, San Diego, CA) at the Joint Genome Institute to generate 51 bp single-end reads (Supplemental Table 2).

Pre-processing of sRNA-seq data were performed as previously described (Stacey et al., 2016). Data were incorporated into an sRNA database and are available for viewing online at https://mpss.danforthcenter.org/dbs/index.php?SITE=maize_sRNA4. Briefly, we first used Trimmomatic version 0.32 to remove the linker adaptor sequences (Bolger et al., 2014). The trimmed reads were then mapped to version 4 of the B73 maize genome (Jiao et al., 2017) using Bowtie (Langmead et al., 2009) allowing zero mismatches, and reads of length 18-34-nt retained. Reads mapping to structural RNAs were then removed and counts were then scaled by multimapping rate, e.g., a read mapping to two locations receives a count of 0.5 at each location. Read abundance was then normalized to library size by scaling to Counts Per 5 Million mapped reads (CP5M) to allow for direct comparison across libraries. For the whole genome analysis, unmapped reads and reads mapping to greater than 50 locations were excluded from further analysis; for the 20 Mb regional analysis, there was no upper limit on multi-mapping rate. Counts were then split into the principal small RNA size classes −21nt, 22nt and 24 nt - and counts summarised into 100 bp fixed windows, tiling the genome.

One sample had less than 1 million mapped reads (PH207 x B73 F_1_ leaf sample) and was omitted from further analysis.

### mRNA-seq library construction and sequencing

RNA-seq samples are as described in (Li et al., 2019) and were downloaded from SRA. The samples analysed in this study, which were paired with the sRNA samples, are described in Supplemental Table S3. Briefly, as detailed in (Li et al., 2019) total RNA was extracted using the miRNAeasy Mini Kit (Qiagen). Extracted RNA was DNase treated using the TURBO DNA-free kit (Life Technologies). Sequence libraries were prepared by the Joint Genome Institute following the standard TruSeq Stranded mRNA HT library preparation protocol (Illumina, San Diego, CA). Samples were sequenced on an Illumina HiSeq 2500 (Illumina, San Diego, CA) at the Joint Genome Institute to generate 150bp paired-end reads. For each RNA-seq library 21-52 million reads were sequenced.

Reads were trimmed using Trimmomatic (Bolger et al., 2014) and mapped to the B73v4 genome (Jiao et al., 2017) by alignment software STAR (Dobin et al., 2013). Uniquely mapped reads were assigned to and counted for the 46,117 B73v4 gene models using FeatureCounts (Liao et al., 2014). Raw read counts were then normalized by library size and accounted for the effect of extremely differentially expressed genes using the TMM (trimmed mean of M values) normalization approach to give CPMs (Counts Per Million reads) for each gene model (Robinson and Oshlack, 2010).

### Annotation of genomic features

The background genomic distribution of genomic features was determined by annotating each 100 bp tile of the genome; as either non-coding (e.g. miRNA and lncRNA), genic, gene-proximal (within 2kb of a gene), transposable element (TE) or intergenic using the genome reference B73 RefGen_v4 and annotation based on Gramene version 36 and miRbase release 22. Synteny classifications (i.e., syntenic and non-syntenic) and assignment to maize sub-genomes were obtained from a previous study based on pairwise whole-genome alignment between maize and sorghum, downloaded from Figshare Schnable 2019: DOI:10.6084/m9.figshare.7926674.v1 (Schnable et al., 2011). The B73 transposable element annotation was sourced from (Anderson et al., 2019).

### Statistical analysis

PCA analysis was performed using the r package pcaMethods. Counts were log2 transformed, scaled by unit variance and clustered using singular value decomposition (flags: scale = “uv”, center = T, method = “svd”).

### Accessions numbers

sRNA data generated in this study is available in the SRA under accession SRA793603 or JGI Proposal ID 1810. The specific SRR numbers for the samples analysed are listed in Supplemental Table 2. The sRNA data and a genome browser are also available for viewing online at https://mpss.danforthcenter.org/dbs/index.php?SITE=maize_sRNA4. RNA-seq data used in this study was previously reported in (Li et al., 2019) and the SRR numbers for the samples analysed in this study are listed in Supplemental Table 3.

## Supplemental Material

Supplemental Table S1 – Overview of the crossing scheme for the maize hybrids analysed in this study.

Supplemental Table S2 – Summary of the sRNA-seq samples generated in this study.

Supplemental Table S3 – Summary of the RNA-seq samples analysed in this study.

## Acknowledgements

DOE supported this work via JGI and GLBRC. PAC and NMS were supported by a grant from NSF (IOS-1802848). RH and BCM were supported by a grant from NSF (IOS-1754097). We acknowledge the assistance of Katie Heslip, MSU with RNA extracts. Computational support was provided by the Minnesota Supercomputing Institute.

---

**Figure S1.**
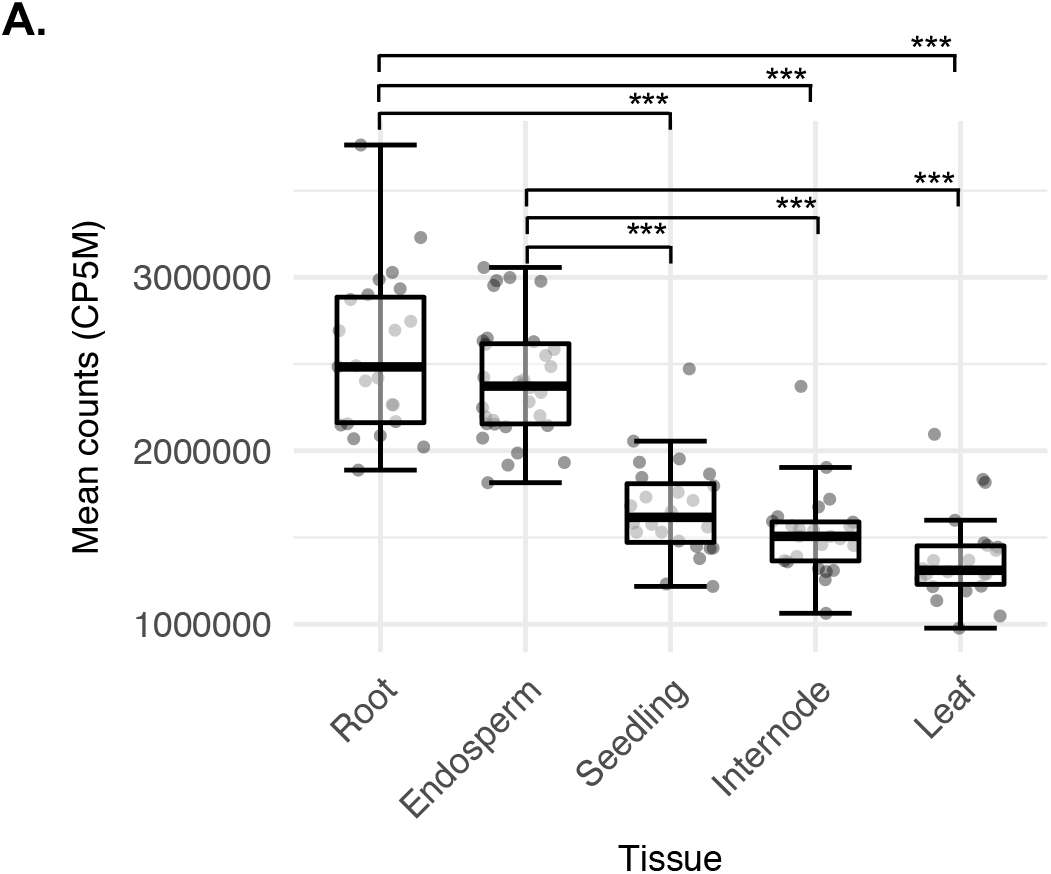
Comparison of the abundance of highly multi-mapping sRNAs between tissues **A** sRNAs mapping to greater than 50 locations in the genome. ANOVA, Tukey’s HSD ***p < 0.001; seedling-root n=20, leaf n=18, seedling-shoot n=21, endosperm n=28, internode n=21; whiskers correspond to the first and third quartiles (the 25th and 75th percentiles).

**Figure S2.**
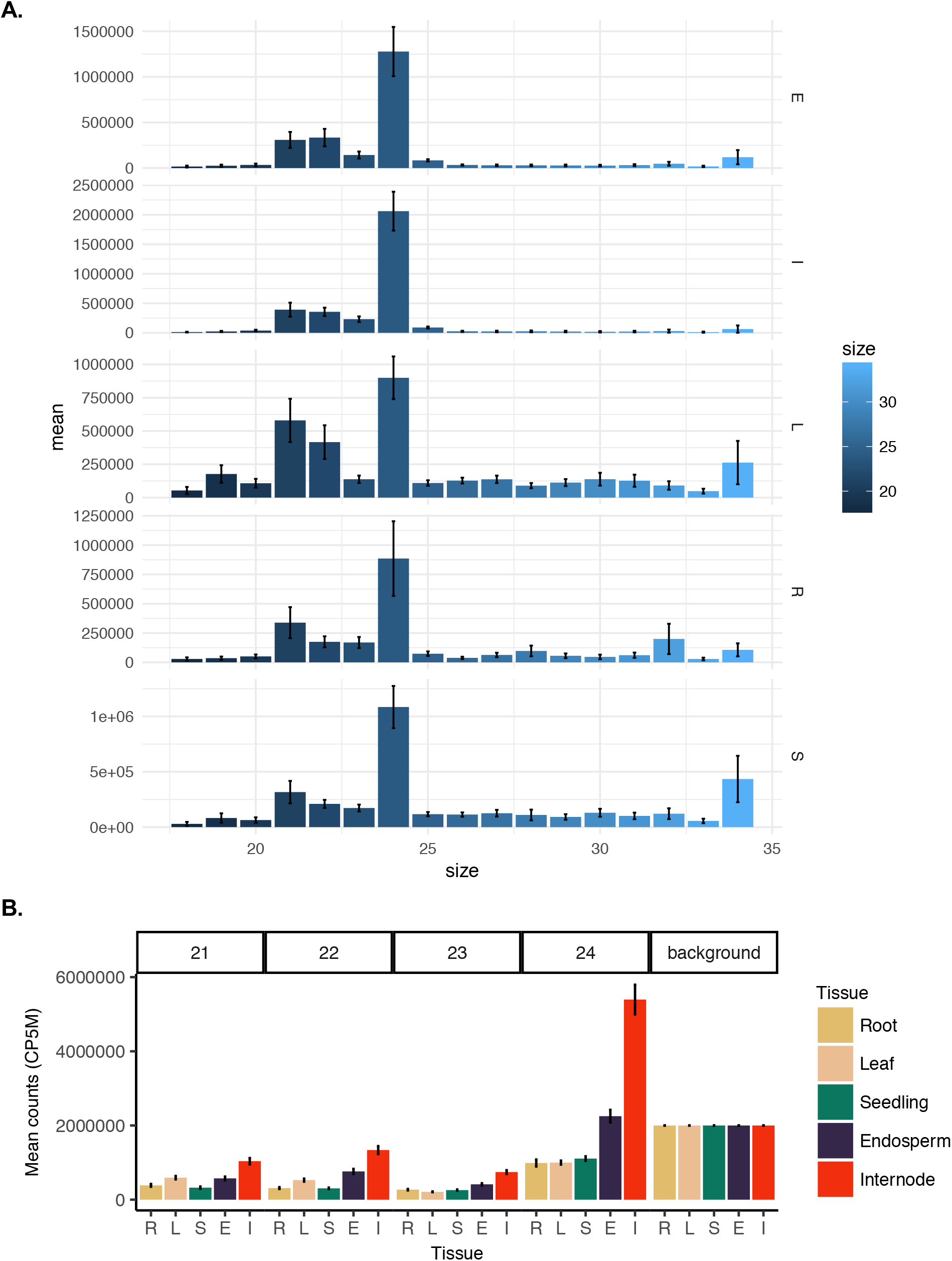
abundance of sRNA size classes. **A.** Average sRNA counts for each size class (18-34-nt) among genotype. **B.** Normalised sRNA abundance. Small RNA abundance was normalised to “background” reads abundance using the abundance of 25-34-nt sRNAs. Error bars denote standard error among genotypes. R = “seedling-root” (n=20), L = “leaf” (n=18), S = “seedling-shoot” (n=21), E = “endosperm” (n=28), I = “internode” (n=21).

**Figure S3.**
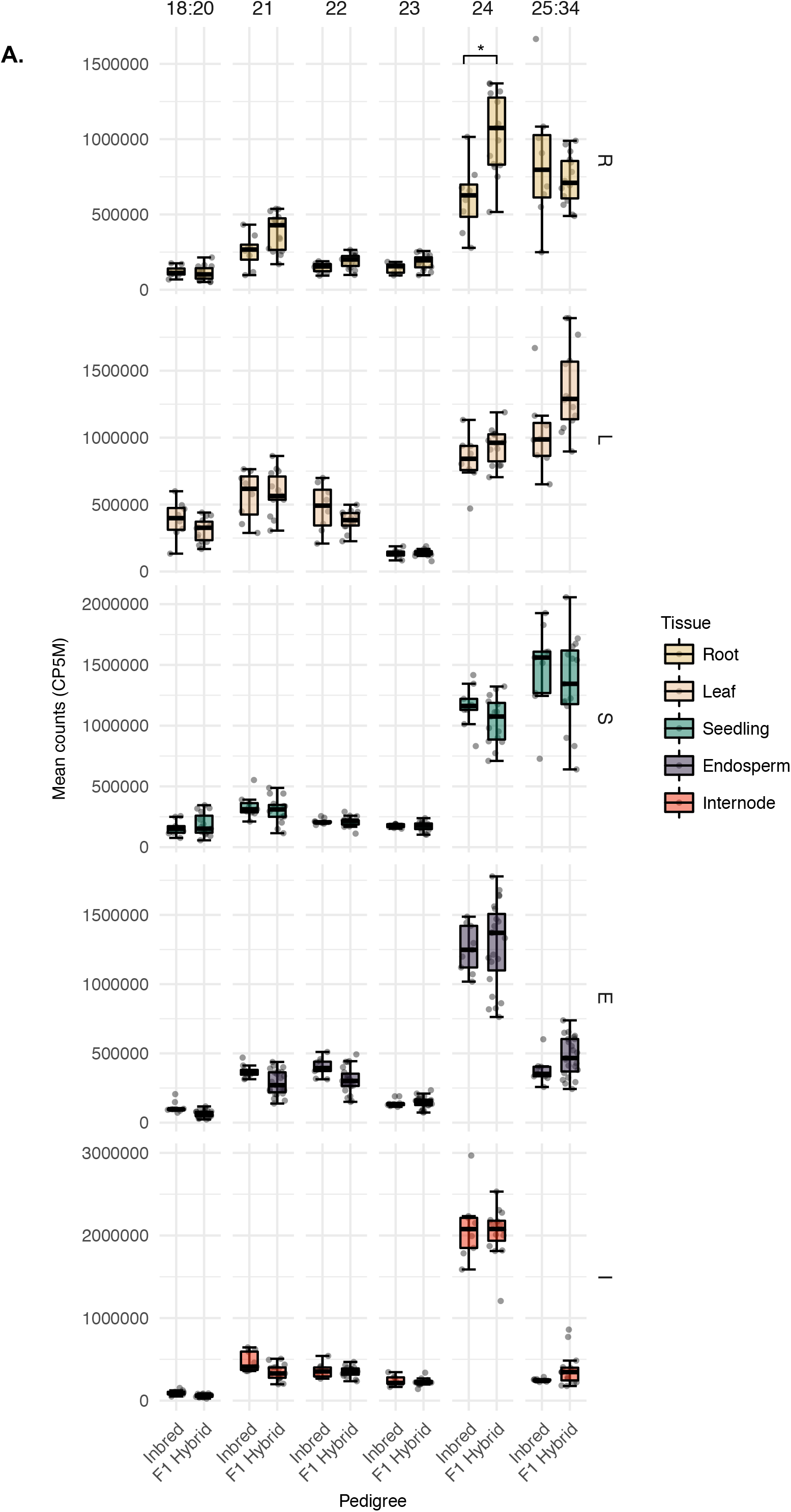
A comparison of sRNA abundance between inbreds and hybrids. **A.** comparison of sRNA abundance between inbreds and hybrids for each tissue. Each dot represents a genotype, whiskers correspond to the first and third quartiles (the 25th and 75th percentiles). R = “seedling-root” (n=20), L = “leaf” (n=18), S = “seedling-shoot” (n=21), E = “endosperm” (n=28), I = “internode” (n=21). ANOVA, Tukey’s HSD * p < 0.05.

**Figure S4.**
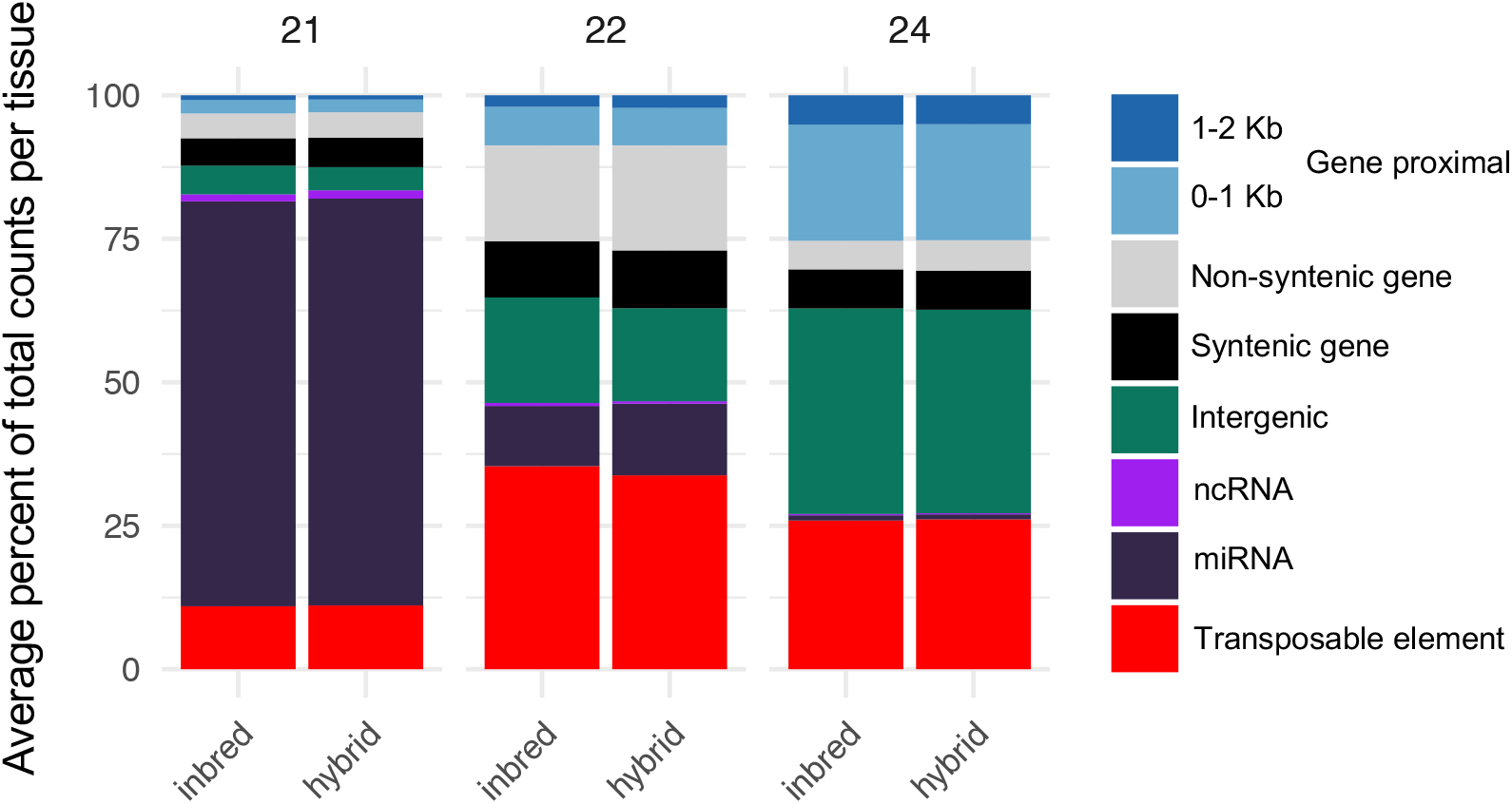
genomic features of sRNA loci. The distribution of genomic features was determined by annotating each 100 bp tile of the genome as either non-coding RNA (e.g. miRNA and lncRNA), genic, gene-proximal (within 2kb of a gene), transposable element (TE) or intergenic using the genome reference B73 RefGen_v4 and annotation based on Gramene version 36 and miRbase release 22. Each meta-feature was then divided into constituent features and the average distribution of sRNA expression determined per tissue. There is no statistically significant difference between inbreds and hybrids for any feature in any size class (t-test, Benjamini & Hochberg adj p value < 0.01).

**Figure S5.**
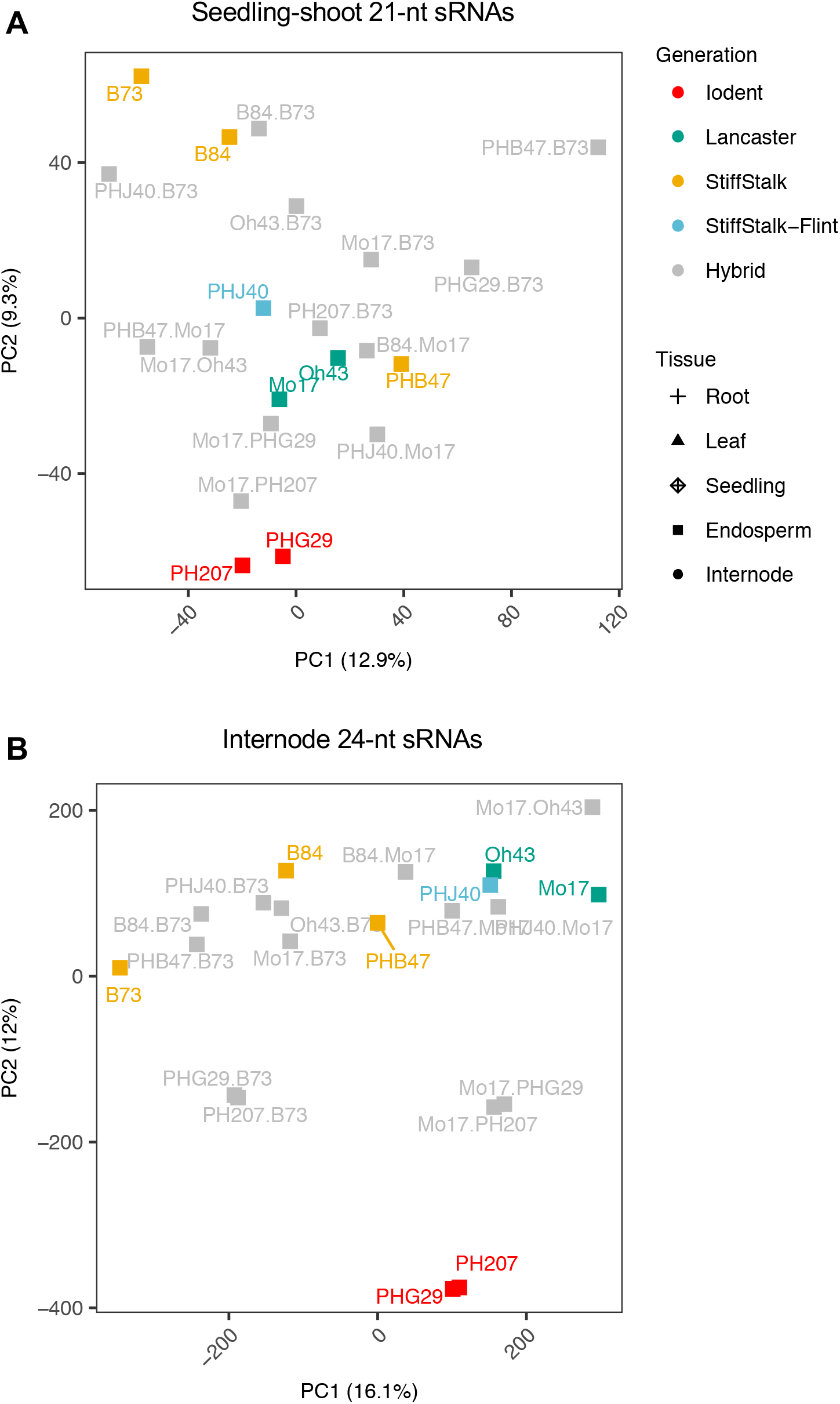
PCA on expressed loci for. Principle component analysis (PCA) on expressed loci for **A.** seedling-shoot 21-nt sRNAs, and **B.** internode 24-nt siRNA. For each sRNA size class PCA was performed using expressed loci with greater than 10 CP5M in at least one sample. Counts were log2 transformed, scaled by unit variance and clustered using singular value decomposition. Colours represent genotype (heterotic group or hybrid), symbols represent tissue type. Percentages in brackets refer to the percent of variance explained by each PC.

**Figure S6.**
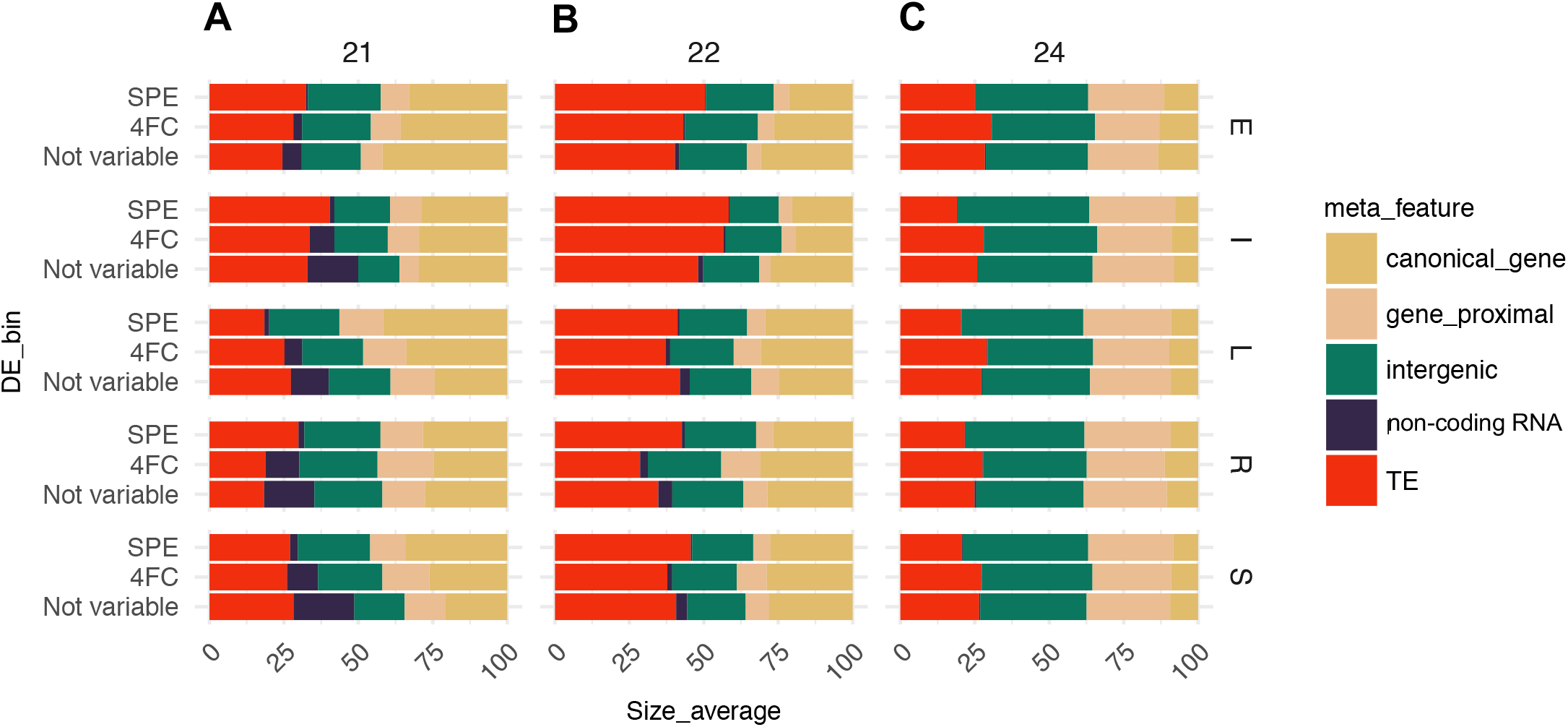
Source of expression variation between inbreds per tissue. **A-C.** Genomic loci contributing variable expression of sRNAs. The distribution of sRNA expression (CP5M) across meta-features for each sample was averaged per tissue and per variable expression category to determine the meta-feature distribution for each size class. For each hybrid the variability in expression between the parents was assessed. Only loci expressed in at least one of the parents were considered (10 CP5M or greater in one parent) by calculating the log2 ratio of high parent/low parent. The proportion of loci in each variable expression category (“not variable”, “2-4 fold”, “> 4 fold” and “SPE”) was collated and then averaged per tissue and size class combination. SPE strictly defined as > 10 CP5M in one parent and 0 in the other parent. R = “seedling-root” (n=13 contrasts), L = “leaf” (n=11 contrasts), S = “seedling-shoot” (n=13 contrasts), E = “endosperm” (n=20 contrasts), I = “internode” (n=13 contrasts).

**Figure S7.**
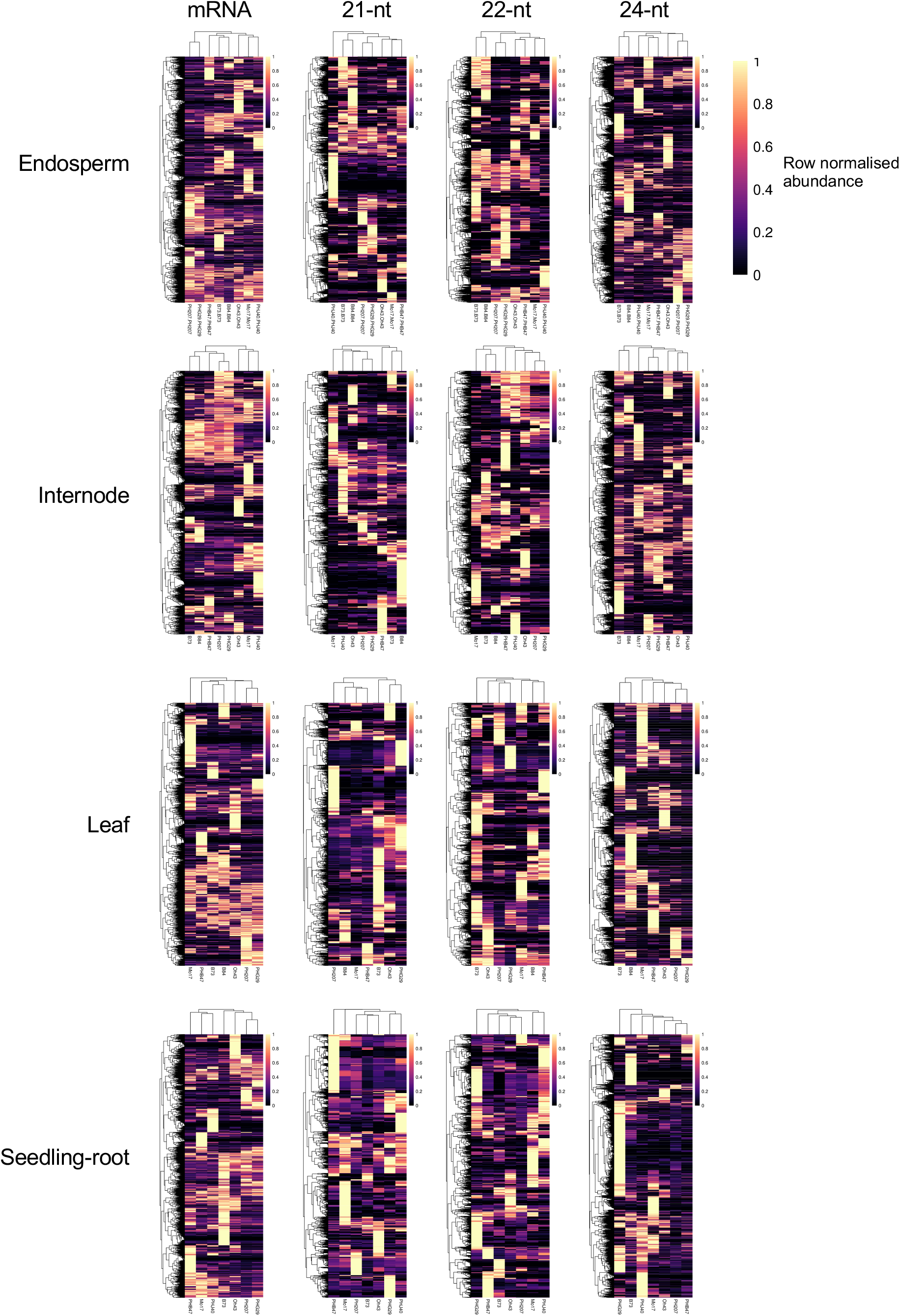
Expression patterns of loci with variable expression in each tissue. Per size class and tissue, each loci with variable expression (>4-fold or SPE) in at least one contrast was profiled for each hybrid. Expression was normalised to the maximum expression (0-1 scale) for that loci and hierarchically clustered. For visualisation a maximum of 45,000 were included per heatmap, loci were randomly subsampled down to 40,000 using the seed 27 for reproducibility.

**Figure S8.**
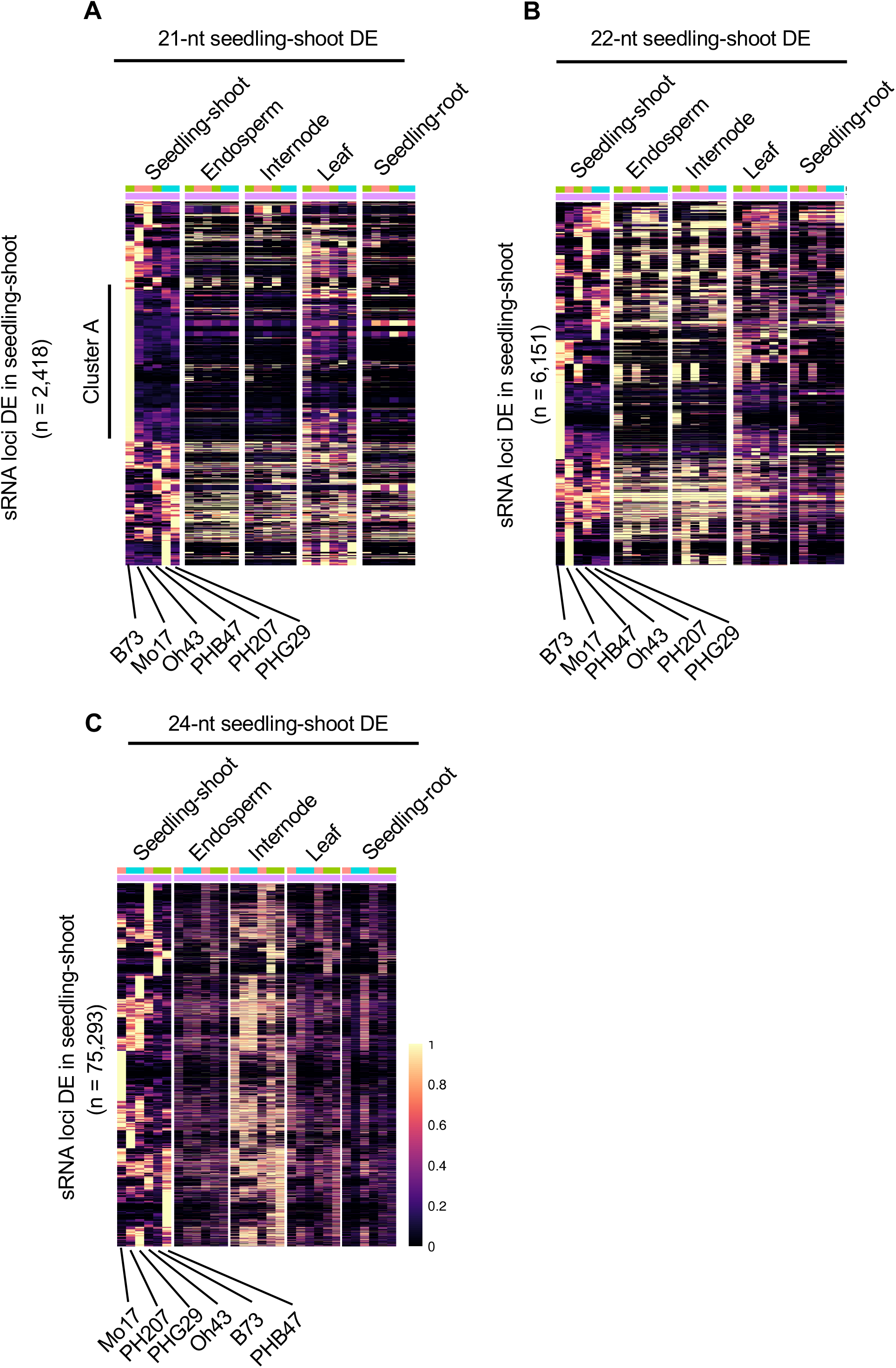
Comparison of variable loci across tissues. For each size class, loci differentially expressed (> 4-fold or SPE) in seedling-shoot were clustered by row and column and the hierarchical trees extracted. Using the list of loci ordered by the hierarchical trees, expression values for the other tissues were extracted for these seedling-shoot DE loci and plotted according to the original clustering in shoot. Each row is normalised to the maximum value in seedling-shoot, with an upper limit of 1 (even if expression is higher in another tissue). The number of variable loci per size class is shown in parentheses, for visualisation a maximum of 40,000 were included per heatmap. This analysis was limited to the six inbreds with complete data in all tissues, PHJ40 and B84 excluded.

**Figure S9.**
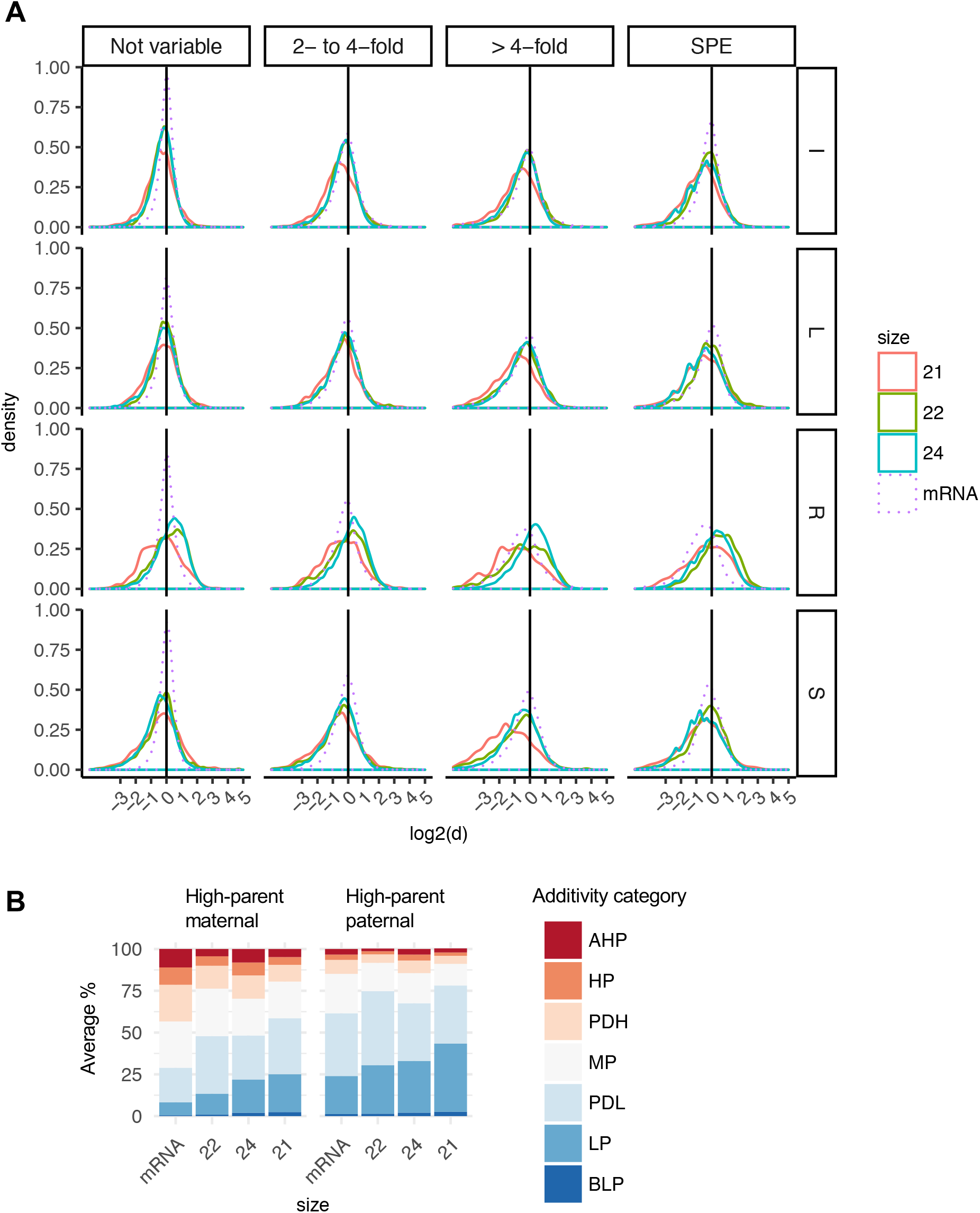
Additivity across tissue and size classes. **A.**Tissue specific analysis of the distribution of additivity in hybrids per sRNA size class compared to mRNA. Additivity was calculated as log2(hybrid / mid-parent) for sRNA and mRNA loci with at least 10 CP5M (2 CPM for mRNA) in one parent and divided into categories for the level of expression variation between the inbred parents. SPE = single parent expression. For plotting the tails were concatenated at +/− 5. L = “leaf”, I = “internode”, R = “seedling-root”, S = “seedling-shoot”. **B** Distribution of additivity categories for sRNA loci that are 4-fold or greater variable between inbred parents in endosperm. Loci are divided by whether the maternal or paternal parent has high expression. The degree of dominance (d/a) calculated as [hybrid – mid-parent/(high-parent – low-parent/2)] was determined. d/a values were then divided into either additive mid-parent or 6 non-additivity levels: Above High Parent (AHP) > 1.25; High Parent (HP) 0.75: 1.25; Partial Dominance High Parent (PDH) 0.25: 0.75; Mid-parent (MP) −0.25: 0.25; Partial Dominance Low Parent (PDL) −0.75: 0.25; Low Parent (LP) −0.75: −1.25; Below Low Parent (BLP) < −1.25.

**Figure S10.**
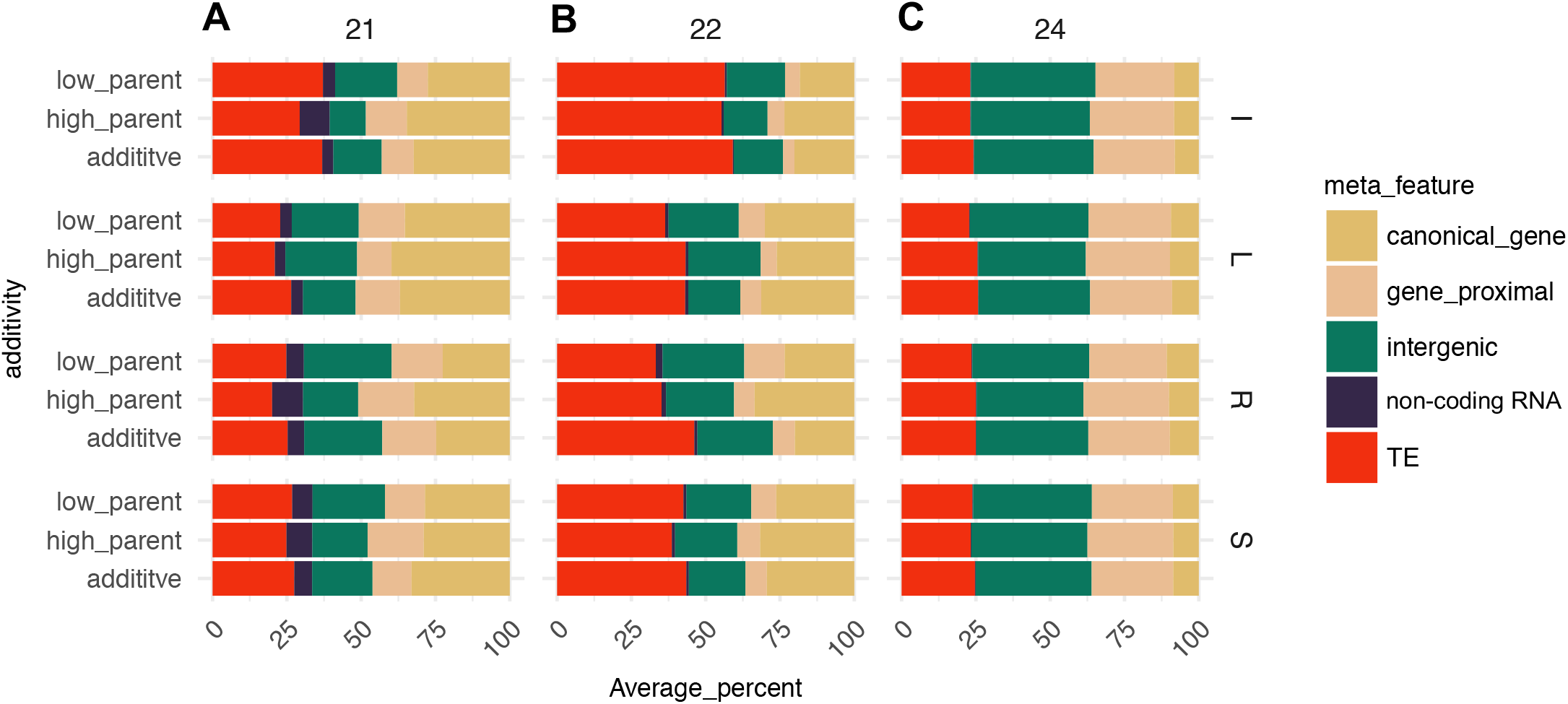
Distribution of genomic features between additive and non-additive loci across tissues. **A-C.** For additive (MP) non-additive high-parent like (AHP, HP and PHD) and non-additive low-parent like (PDL, LP, BLP), the distribution of sRNA expression (CP5M) across meta-features for each sample was averaged per tissue for each size class. R = “seedling-root” (n=8 contrasts), L = “leaf” (n=7 contrasts), S = “seedling-shoot” (n=7 contrasts), I = “internode” (n=8 contrasts). Endosperm was excluded owing to the genome imbalance.

